# Brain Network Architecture Constrains Age-related Cortical Thinning

**DOI:** 10.1101/2022.07.08.499363

**Authors:** Marvin Petersen, Felix L. Nägele, Carola Mayer, Maximilian Schell, D. Leander Rimmele, Elina Petersen, Simone Kühn, Jürgen Gallinat, Uta Hanning, Jens Fiehler, Raphael Twerenbold, Christian Gerloff, Götz Thomalla, Bastian Cheng

## Abstract

Age-related cortical atrophy, approximated by cortical thickness measurements from magnetic resonance imaging, follows a characteristic pattern over the lifespan. Although its determinants remain unknown, mounting evidence demonstrates correspondence between the connectivity profiles of structural and functional brain networks and cortical atrophy in health and neurological disease. Here, we performed a cross-sectional multimodal neuroimaging analysis of 2633 individuals from a large population-based cohort to characterize the association between age-related differences in cortical thickness and functional as well as structural brain network topology. We identified a widespread pattern of age-related cortical thickness differences including “hotspots” of strong age effects located in brain areas with high centrality (structural network hubs). Regional age-related differences were furthermore strongly correlated within the structurally defined node neighborhood. The overall pattern of thickness differences as well as its change throughout the later lifespan was found to be anchored in the functional network hierarchy as encoded by macroscale functional connectivity gradients. Lastly, the identified difference pattern covaried significantly with cognitive and motor performance. Our findings indicate that connectivity profiles of functional and structural brain networks might act as organizing principles behind age-related cortical thinning as an imaging surrogate of cortical atrophy.

## Introduction

Understanding the neurobiological processes underlying aging is a critical challenge given the increasing average age of societies and growing incidence of age-related neurological impairments worldwide (Beard et al., 2016). Over the lifespan, changes in brain structure and function accrue, leading to reduced performances in multiple cognitive and motor domains like executive function, memory, attention as well as general muscle strength and mobility (Baciu et al., 2015; López-Otín et al., 2013; Tromp et al., 2015). Collectively, these changes affect well-being and psychosocial functioning and independence in advancing age.

Magnetic resonance imaging (MRI) provides an avenue to investigate changes in brain structure and function during aging in vivo. MRI-based reconstruction of cortical morphology enables the assessment of cortical thickness. Cortical thinning is established as a valuable imaging marker of age-related grey matter atrophy linked to decline in cognitive and motor functions (Clark and Taylor, 2011; Pacheco et al., 2015). As demonstrated by previous epidemiological studies, age-related thinning of the cerebral cortex follows a nonuniform trajectory, the determinants of which are not well understood (Frangou et al., 2021). Cortical thickness reaches its peak within the first decade of life, followed by a dynamic trajectory of thinning starting off with a steeper decline during the first three life decades, which finally decelerates over the remaining lifespan (Frangou et al., 2021; Walhovd et al., 2017). Intriguingly, age-dependent thinning of the cerebral cortex does not occur in a homogeneous pattern: initially occurring primarily in association cortices, foci of cortical thinning appear to shift towards primary sensorimotor areas during later stages of aging (Appleton et al., 2020; Davis et al., 2009; Douaud et al., 2014; Frangou et al., 2021).

Brain network analysis, commonly referred to as “connectomics”, has proven insightful in elucidating the underlying mechanisms of age-related cortical thinning (Fornito et al., 2015; Fornito and Bullmore, 2015). Previous reports indicate that during late adulthood, increased cortical thinning preferentially occurs in regions connected by white matter tracts that demonstrate increased age-related structural disintegration (Storsve et al., 2016). Moreover, resting-state functional connectivity changes have been shown to co-occur with cortical thinning (Schulz et al., 2022; Vieira et al., 2020). Although there is evidence for interactions between brain network connectivity (i. e., the connectome) and cortical morphology in general, investigations of age-related cortical thinning in association with connectome topology are scarce.

Amassing evidence from joint MRI and clinical investigations demonstrates a strong link between disease-related alterations of the cerebral cortex and connectome topology: for example, brain areas with prominent cortical atrophy in primarily neurodegenerative forms of dementia appear to be strongly structurally and functionally connected (Savard et al., 2022; Seeley et al., 2009). Moreover, these conditions appear to preferentially affect brain areas located at the associative-transmodal regions, highlighting the relevance of the functional network hierarchy in pertaining pathomechanisms (Greicius et al., 2004; Hu et al., 2022). In patients with schizophrenia, cortical thinning is primarily observed in “neighborhoods” of functionally and structurally highly interconnected brain regions (Shafiei et al., 2020). Subcortical stroke induces cortical thinning in remote, yet connected brain regions (Cheng et al., 2019; Mayer et al., 2020). Lastly, network hubs – i.e, nodes featuring high connectivity and prominent location within a network – are preferentially targeted in manifold diseases due to their topological centrality and high metabolic demands (Crossley et al., 2014).

Based on this evidence, we hypothesized that the pattern of age-related interindividual cortical thickness differences – as a cross-sectional proxy of age-related cortical thinning – is associated with principal aspects of functional and structural connectome topology. To address this hypothesis, we assessed if the effect of age on cortical thickness differences occurs (1) preferentially in network hubs; (2) in highly interconnected network neighborhoods and (3) if the age-related pattern of cortical thickness differences follows the constraints imposed by the functional network hierarchy as encoded in macroscale functional connectivity gradients (Margulies et al., 2016). For this purpose, we contextualized the pattern of age-related cortical thickness differences and connectome measures in a surface-based spatial correlation analysis in MRI data from participants of a large-scale, single-center, population-based cohort study (Hamburg City Health Study) (Jagodzinski et al., 2019). Supplementing our analysis of imaging data, we characterized the association between age-related cortical thickness differences and clinical phenotypes, specifically cognitive and motor functions. With this work we aimed to contribute to the undesrstanding of the fundamental principles underlying age-related structural brain changes and their clinical phenotypes.

## Results

### Sample characteristics

Quality checked data from 2633 subjects were included in this analysis. *Table 1* provides an overview of demographic and phenotypical data. Median age was 65 years (IQR=14), 44% of participants were female and median years of education were 13 (IQR=4). Median cortical thickness was 2.5 mm (IQR=0.13).

**Table 1:** Descriptive statistics

|  | Median (IQR) /<br>percentage<br>n=2633 |
| --- | --- |
| Age (years) | 65 (14) |
| Sex at birth (% female) | 0.44 |
| Years of education | 13 (4) |
| TMT A score (sec) | 37 (17) |
| TMT B score (sec) | 82 (42) |
| Animal Naming Test score | 24 (9) |
| Wortliste Recall Sum | 8 (2) |
| Mini Mental State Exam | 28 (2) |
| Geriatric Depression Scale | 1 (3) |
| PHQ9 (% medium or severe depressive symptoms) | 0.26 |
| Hand Grip Strength R (kg) | 34.4 (17) |
| Hand Grip Strength L (kg) | 32.35 (16.36) |
| Time Timed Up And Go Test (sec) | 7 (2) |
| Chair Rising Test (% passed) | 0.97 |
| Tandem Test (% passed) | 0.95 |
| Cortical thickness (mm) | 2.5 (0.13) |

### Age-related interindividual cortical thickness differences

Cortical thickness and age across individuals were related in an adjusted general linear model, revealing a widespread pattern of lower cortical thickness with advancing age as represented by negative *β* estimates (*figure 1*). This effect was strongest within primary somatosensory and motor cortices as well as the superior temporal lobe. Linear relationships were neutral to positive in the anterior and posterior cingulate cortex as well as the inferior temporal lobe.

**Figure 1:**
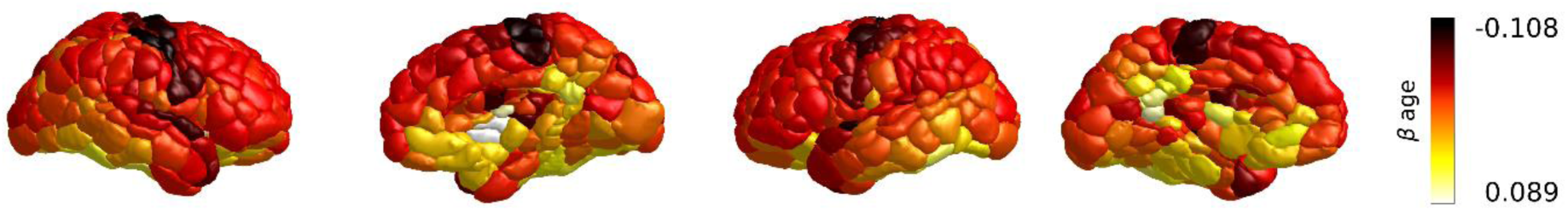
Age-related cortical thickness differences across all individuals. Negative *β* values correspond to lower cortical thickness with higher age, whereas positive *β* values denote higher cortical thickness. Negative *β* values were most pronounced in the primary motor and sensory cortices and the superior temporal lobe. The anterior cingulate cortex, precuneus and inferior temporal lobe showed a neutral to positive linear association between cortical thickness and age. All nodes showed a statistically significant relationship between age and cortical thickness (*p_FDR_* < 0.05). Abbreviations: *p_FDR_*-p-value corrected for false-discovery rate.

### Spatial contextualization of age-related cortical thinning

We investigated patterns of age-related cortical thickness differences in relation to three network topological concepts: hub ranks, neighborhood thickness alteration and macroscale functional connectivity gradients. Therefore, surface-based spatial correlations were performed between *β* estimates and measures of structural and functional connectivity derived from group-level connectomes of the Human Connectome Project Young Adult dataset (*figure 2*).

**Figure 2:**
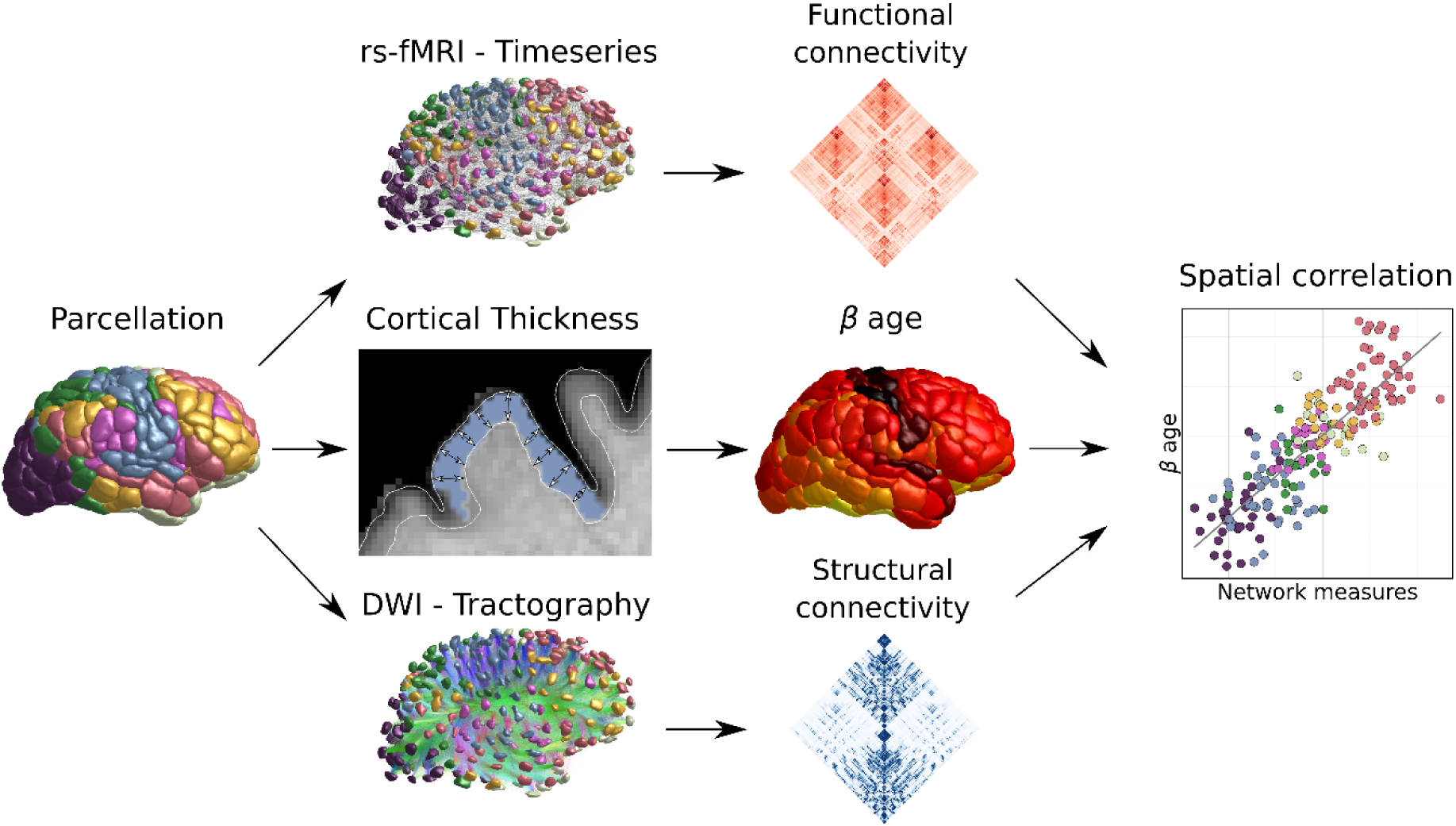
Methodological approach for contextualization of age-related cortical thickness differences (*β* age: nodewise standardized estimate from the general linear model relating age and cortical thickness). See methods section for detailed descriptions. Abbreviations: rs-fMRI - resting-state functional magnetic resonance imaging.

### Hub ranks

Nodes with highest hub ranks in functional connectivity (i.e. nodes with a ranking <100) were identified as sensorimotor and visual cortices. Highest ranking hub nodes based on structural connectivity were superior-frontal, cingulate, sensorimotor and visual areas (node metric results can be found in *supplementary table S1*). Age-related cortical thickness differences showed a significant positive correlation with hub ranks based on both functional (*r_sp_* =0.27, *p_spin_* =0.029, *figure 3a*) and structural connectivity (*r_sp_* =0.22, *p_spin_* =0.047, *figure 3b*). Specifically, cortical areas identified as high-ranking hub nodes demonstrated a lower cortical thickness at higher age compared to lower ranking nodes.

**Figure 3:**
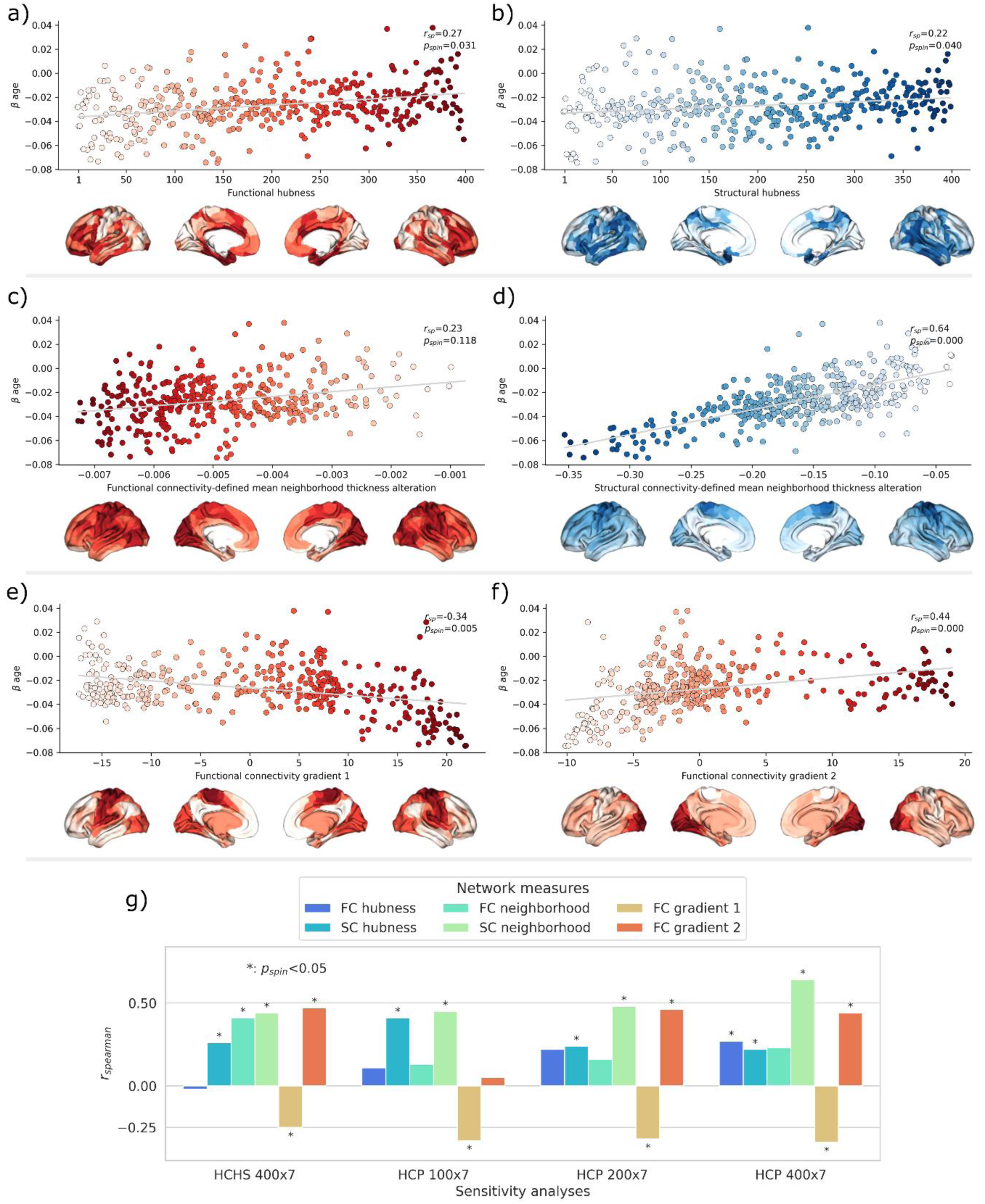
Spatial correlations of age-related cortical thickness differences (*β* age) and brain network connectivity measures (red: structural connectivity, blue: functional connectivity). Scatter plots displaying the spatial relationship are supplemented by surface maps for anatomical localization. a) structural and b) functional hubness rank. Lighter colors represent a higher hubness ranking. c) structural and d) functional neighborhood thickness alteration. e) & f) Intrinsic functional network hierarchy represented by functional connectivity gradient scores 1 and 2. g) Results of sensitivity analyses. Correlation results and significance are demonstrated for analysis pipelines including group-averaged Schaefer400x7 connectomes from HCHS subjects as well as Schaefer100x7, Schaefer200x7 and Schaefer400x7 connectomes from HCP subjects. Abbreviations: *β* age - *β* estimate describing the linear relationship between age and cortical thickness; *r_sp_* - Spearman correlation coefficient; *p_spin_* - p-value derived from spin permutation results.

#### Neighborhood thickness alteration

Nodes identified in primary sensorimotor brain areas showed most pronounced neighborhood thickness alteration in terms of lower cortical thickness based on neighborhoods defined by structural connectivity. Nodes identified in cingulate and parietal brain areas remained relatively unchanged (*supplementary table S1*). A positive and significant association (*r_sp_* =0.64, *p_spin_* <0.001) of neighborhood thickness alteration with age-related cortical thickness differences was found (figure 3d), indicating lower cortical thickness at higher age in brain areas highly connected to other brain areas exhibiting similar age-effects.

Primary somatosensory and visual nodes showed most pronounced neighborhood thickness alteration in terms of lower cortical thickness based on functional connectivity measures. A positive, non-significant (*r_sp_* =0.23, *p_spin_* =0.118) association with age-related cortical thickness differences was found (figure 3c).

#### Macroscale functional connectivity gradients

Nodes were embedded in the intrinsic functional cortical hierarchy along the first and second macroscale functional connectivity gradients (*figure 3e and 3f*, brain surfaces): Gradient 1 spanned the sensorimotor-associative axis whereas the ends of gradient 2 were anchored in sensorimotor as well as visual cortices, analogous to previous reports (Margulies et al., 2016). A scree plot of the respective eigenvalues is shown in the supplementary materials (*supplementary figure S2*). Scores of both, the first and second functional connectivity gradient, were significantly spatially correlated with the pattern of age-related cortical thickness differences (*r_sp_* =-0.34, *p_spin_* =0.005, *figure 3e*; *r_sp_* =0.44, *p_spin_* <0.001, *figure 3f*). This indicates that the differential degree of age-related differences within our sample is captured by both functional connectivity gradients. In fact, brain regions of lower cortical thickness at higher age were located in primary and unimodal sensorimotor regions (*figure 4a*). The top 10 percent of nodes with the most negative *β* values had median gradient 1 and 2 scores of 18.44 and -7.88, respectively (*figure 4a*, red arrow). On the other end of the distribution of *β* values, brain regions of higher cortical thickness at higher age were located in associative-transmodal regions. Top 10 percent of nodes with most positive *β* values had median gradient 1 and 2 scores of -5.99 and -0.77 respectively (see *figure 4a, blue arrow*).

**Figure 4:**
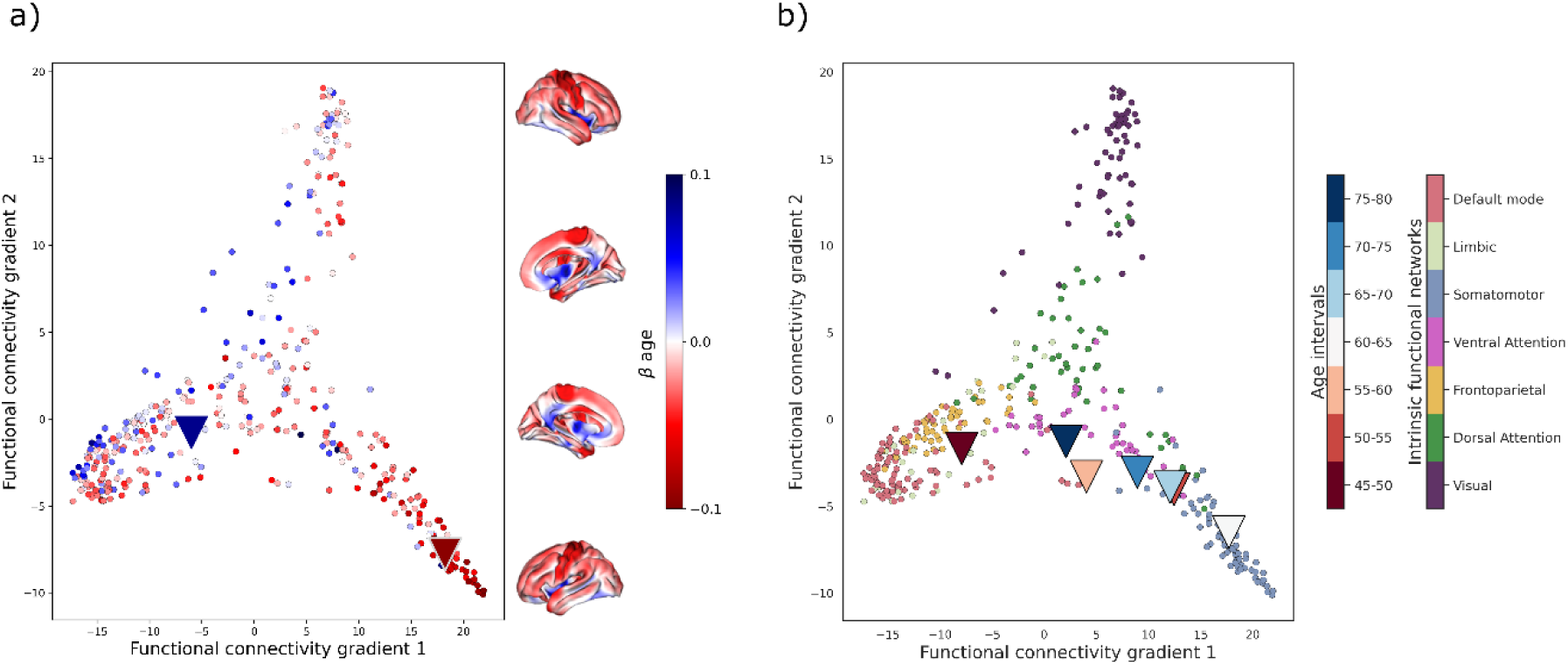
Age effect foci localized in gradient space. Nodes represented by dots are plotted according to their location in gradient space as indicated by their functional connectivity gradient scores. Here, visual areas are located at the top, associative areas at the lower left and sensorimotor areas at the bottom right. a) Age effect foci in all individuals. Nodes are color-coded by their respective linear relationship between age and cortical thickness (*β*), where negative values indicate a lower cortical thickness at higher age. *β* values are also plotted on the cortical surface for anatomical localization. Colored arrow heads indicate median gradient scores of the nodes with 10 percent of strongest negative (red) and strongest positive (blue) *β*-values. b) Negative age effect foci grouped by age intervals. Negative age effect foci were defined as the nodes with the 10 percent most negative *β*- values. Arrows denote median gradient scores of negative age effect foci and are colored by age intervals. Nodes are colored by their membership to intrinsic functional networks according to the Schaefer atlas.

Analysis by age interval subgroups provided information about localization of age effect foci in younger and elderly individuals in relation to both gradients (figure 4b). In younger individuals (age 45-50 years), the negative relationship between age and cortical thickness was steepest in associative-transmodal brain regions constituting the limbic, default mode and frontoparietal intrinsic functional networks. In individuals with advancing age intervals (50 to 65 years), age-effects on cortical thickness were most pronounced in unimodal sensorimotor areas. Lastly, for the oldest individuals (>65 years), most negative *β* values were mainly focused on brain regions located centrally within functional connectivity gradient 1, i.e., task-positive networks like the ventral and dorsal attention network.

### Sensitivity Analysis

Across differing cortical parcellation schemes derived from HCHS and HCP connectomes, hub ranks, the structurally-defined neighborhood thickness alteration and the functional connectivity gradient score 1 maintained a significant relationship to age-related cortical thickness differences (*β*) in line with results from our main analysis (figure 3g, *supplementary table S3 and figure S4*). Of note, the second functional connectivity gradient derived from the Schaefer100x7-parcellated HCP-connectome spanned from visual cortices to associative-transmodal instead of sensorimotor cortices, potentially obscuring the spatial correlation of *β* and the gradient 2 score in this particular correlation.

### Association between cortical thickness, age and clinical phenotypes

Partial least squares analysis identified 3 significant latent variables relating age, sex, education, neuropsychological test scores, motor test scores and nodewise cortical thickness measures. The first latent variable explained the majority (85.10%) of the shared covariance and was thus chosen for subsequent analysis (*figure 5a; supplementary table S5*). Specifically, the first latent variable represented a predominant covariance profile consisting of younger age and better performance both in neuropsychological (e.g., shorter times to complete the TMT) as well as motor tests (e.g., shorter timed up and go interval; *figure 5b*). Of note, age showed the strongest contribution among all variables to the covariance profile as indicated by highest loading to the first latent variable.

**Figure 5.**
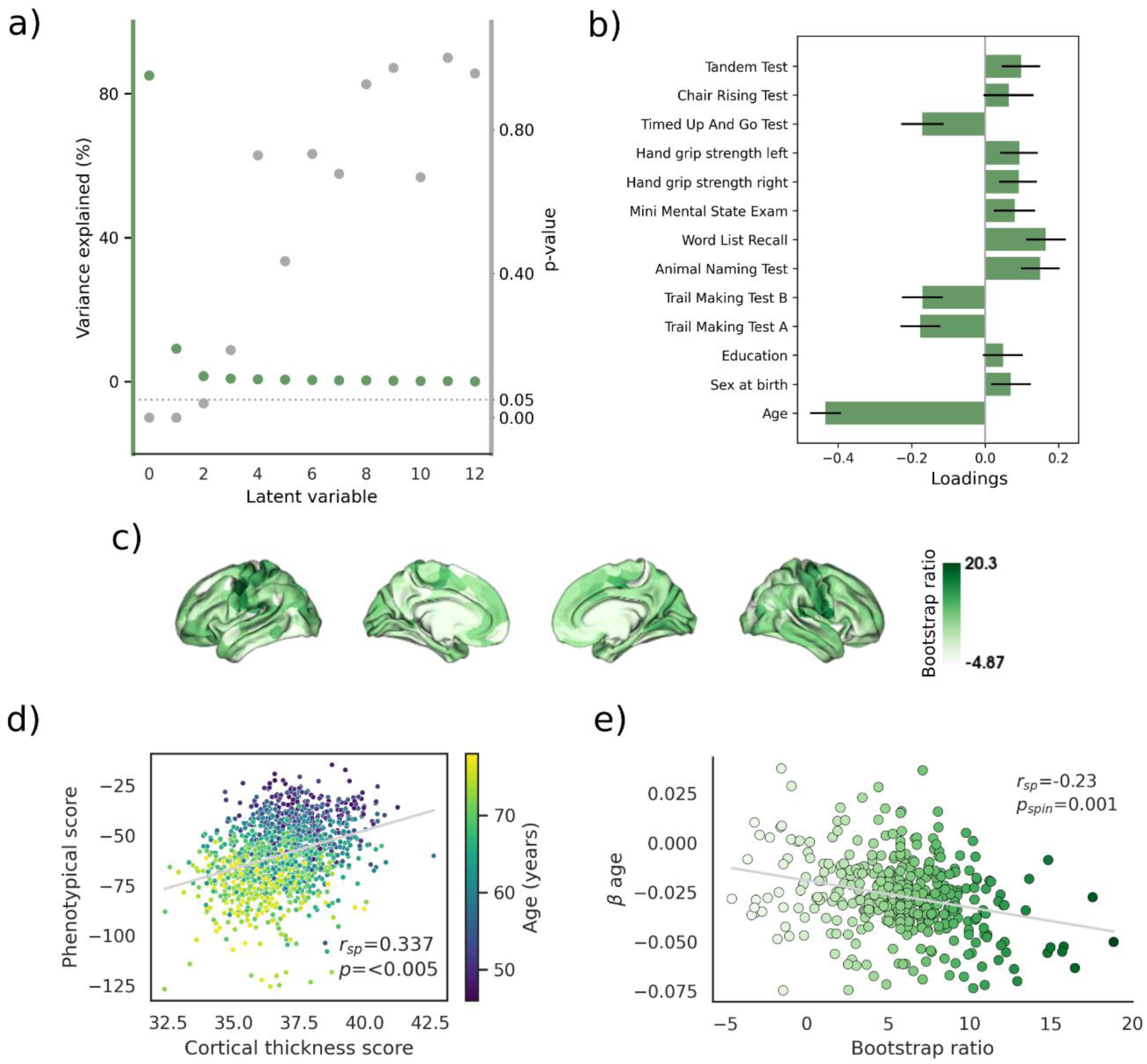
Results from partial least squares analysis. a) Explained variance and significance levels of identified latent variables. The first significant latent variable explaining 85.10% of variance was used for subsequent analysis. b) Phenotypical covariance profile of the first latent variable. c) Covarying cortical thickness pattern represented by the bootstrap ratio with higher ratios indicating larger contribution of brain areas to the overall covariance profile d) Correlation of individual phenotypical as well as cortical thickness scores. Higher scores represent more pronounced expression of the covariance profile as exemplified by age-wise coloring. e) Spatial relationship of the bootstrap ratio to the pattern of age-related cortical thickness differences. In the scatterplot dots represent Schaefer400x7 parcels. Dots are colored by the respective bootstrap ratio.

Bootstrapping was performed to identify nodes that contributed relevantly to the covariance profile of the chosen first latent variable. Cortical thickness in sensorimotor areas contributed highest to the shared covariance of the first latent variable as indicated by a strong positive bootstrap ratio. That indicates that lower cortical thickness in sensorimotor regions corresponds with higher age, worse cognitive and motor test performances and vice versa.

Individual cortical thickness and phenotypical component scores for latent variable 1 were computed. These scores represent the degree by which an individual expresses the corresponding covariance profiles regarding either cortical thickness or phenotype. As per definition, scores were significantly correlated (*r_sp_* =0.34, *p_spin_* <0.005, *figure 5d*) indicating that individuals matching the phenotypical covariance profile also expressed the cortical thickness pattern. The bootstrap ratio was significantly spatially correlated with age-related cortical changes (*r_sp_* =-0.23, *p_spin_* =0.001; *figure 5e*). Thus, a correspondence between age-related cortical thickness differences and the identified covariance profiles could be established visually as well as statistically. Abovementioned results remained stable in a supplementary analysis if age, sex and years of education were excluded from the phenotypical variables (*supplementary figure S6*).

## Discussion

In this work, we link principal organizational aspects of structural and functional brain networks with age-related cortical thickness differences based on MRI data from a large, population-based epidemiological study. We report three main findings: (1) The pattern of age-related cortical thickness differences is conditioned by the brain network architecture, specifically structural connectivity to brain areas with shared age-effects, i.e., neighborhood thickness alteration. (2) Patterns of age-dependent cortical thickness differences correspond well with the intrinsic macroscale cortical organization expressed by functional connectivity gradients. Along these gradients, “hotspots” of negative age effects on cortical thickness were localized in differing brain regions for subgroups of increasing age intervals. (3) Age effects on cortical thickness were strongest in brain regions associated with clinical phenotypes of worse neuropsychological and motor performance.

Relating age and cortical thickness in a general linear model revealed a widespread pattern of negative *β* values which were most pronounced in primary sensorimotor regions (*figure 1*). As there is consensus that cortical thickness decreases with advancing age, we interpret the cross-sectionally-derived negative *β* values formally indicating age-related cortical thickness differences as age-related cortical thinning (Frangou et al., 2021; Raz et al., 1997; Storsve et al., 2014; Walhovd et al., 2017). In doing so, we comply with previous cross-sectional studies of age effects on brain morphometry (Frangou et al., 2021; Lemaitre et al., 2012; Salat et al., 2004; Vieira et al., 2020). The thickness of the anterior cingulate cortex, precuneus and inferior temporal cortex remained relatively preserved during aging. These findings correspond well with previous results from epidemiological, cross-sectional and longitudinal studies (Appleton et al., 2020; Frangou et al., 2021; Salat et al., 2004; Storsve et al., 2014; Wierenga et al., 2022).

From a topological perspective on the connectome, brain areas with high structural centrality – i.e., structural hubs – were associated with increased age-related cortical thinning (*figure 3b*). Hubs are brain network nodes characterized by high general connectivity and consequently high metabolic needs (Alexander-Bloch et al., 2013; Liang et al., 2013; Tomasi et al., 2013; van den Heuvel and Sporns, 2013, 2011). Due to this special configuration, hubs exhibit distinct susceptibility to pathology according to the “nodal stress” hypothesis (Crossley et al., 2014). Aging effects like oxidative stress might be most impactful on metabolically most active nodes (Ionescu-Tucker and Cotman, 2021). As the fabric of long-range neural connections is mainly maintained by hubs, they might be especially prone to remote effects induced by age-related microvascular injury which preferentially affect long-range white matter fiber tracts (Petersen et al., 2022, 2020). Moreover, hubs may also be disproportionately affected by hampered axonal transport mechanisms during aging (Milde et al., 2015). Taken together, our findings are in line with the hypothesis of high-centrality brain areas receiving the largest impact of age-related cortical disintegration. The relatively small effect sizes might, however, indicate a subordinate role.

Previous reports suggest that during late adulthood, cortical thinning preferentially occurs in regions connected by white matter tracts that demonstrate increased age-related structural disintegration (Storsve et al., 2016). We therefore tested whether age-related cortical thinning relates to thickness alterations in the functionally and structurally defined neighborhood. We could establish a substantial correlation between age-related cortical thinning in individual brain areas with the collective age-related cortical thinning of their neighborhood defined by structural connectivity (*figure 3d*). In contrast, this association was not found to be significant for neighborhoods defined by functional connectivity. Therefore, our findings indicate that white matter fiber tracts – as the main histological correlate of structural connectivity assessed by MRI tractography – exceed functional connectivity in constraining age-related neurodegeneration. Collectively, we interpret our findings as evidence for a strong interrelation between a brain regions’ age-dependent morphometric change (i.e., thinning) and its underlying structural connectivity profile.

Multiple mechanisms might explain how structural connectivity determines the observed pattern of cortical atrophy beyond localized cellular aging processes (López-Otín et al., 2013). Similar to patterns of cortical atrophy observed in primary neurodegenerative diseases or acute vascular brain injury, white matter fiber tracts may provide a scaffold for propagating age-related cortical atrophy across the network (Agosta et al., 2015; Cheng et al., 2019). Although speculative, degeneration of cortical neurons might be caused by hampered communication via excitotoxicity, diminished excitation and metabolic stress (Saxena and Caroni, 2011), ultimately resulting in dysfunctionality and structural disintegration of connected brain regions (Feeney and Baron, 1986). In addition, previous longitudinal work has shown that age-related white matter alterations might lead to remote degenerative effects in the connected gray matter areas (Storsve et al., 2016). Thus, the occurrence of age-related cortical thinning in connected neighborhoods might be explained by the degree of white matter dysconnectivity which neighboring nodes typically share due to their shared connectivity profile.

Beyond giving rise to phenomena like propagation of cortical atrophy and dysconnectivity, network connectivity might be a reflection of overarching organizational principles that also determine cortical atrophy during the lifespan. A large body of recent literature demonstrates that a multitude of cortical properties follows a sensorimotor-association axis (Sydnor et al., 2021): extremes of functional network hierarchy represented by the principal functional connectivity gradient, evolutionary hierarchy as denoted by expansion during phylogeny, and anatomical hierarchy encoded in myelination degree span from sensorimotor to associative brain regions (Glasser and Van Essen, 2011; Hill et al., 2010; Margulies et al., 2016). The general organizational paradigm captured by this axis is thought to reflect effective hierarchical information processing in the brain: regions of lower rank - unimodal regions like primary sensory and motor cortices - are involved in externally-oriented tasks like perception, whereas higher-rank areas – associative-transmodal regions like the default mode network - integrate collected information to contribute to internally-focused mental faculties of higher order (Mesulam, 1998).

In our work, we tested the hypothesis that the observed overall pattern of cortical thinning adheres to the functional network hierarchy as encoded in the first and second functional connectivity gradients. The former can be considered as a proxy for the canonical sensorimotor-association axis of brain organization. We successfully recovered the first and second functional connectivity gradients from HCP and HCHS connectomes. Over the whole sample, age-dependent cortical thinning generally followed a sensorimotor-fugal pattern with highest effects located in the primary sensory and motor cortices (*figure 1*). Spatial correlations revealed that thinning gradually changed along both, the first and second functional connectivity gradient (*figures 3g and h*), peaking in sensorimotor areas (*figure 4a*, red arrow). Less pronounced thinning or unchanged thickness values across age were localized to associative-transmodal and visual cortices (*figure 4a*, blue arrow). Anatomical, functional and genomic factors influencing age-related neurodegeneration might be differentially distributed along the sensorimotor-association axis, which could explain the corresponding pattern of age-related cortical thinning. Taken together, our findings underline the fundamental role of functional network hierarchy and consequently the sensorimotor-association axis for cortical atrophy in aging.

Over the lifespan, the sensorimotor-association axis also provides the framework for cortical maturation. Specifically, neural maturation advances from sensorimotor regions to associative cortices. The observation that associative areas develop latest ontogenetically, whereas they appear to degenerate first during aging, is referred to as the “last-in first-out”- hypothesis (Appleton et al., 2020; Davis et al., 2009; Douaud et al., 2014). Implied here is that sensorimotor areas mature earlier during ontogeny than associative areas, whereas they degenerate at a later stage. On that basis we hypothesized that throughout aging, cortical thinning foci gradually change along the functional network hierarchy as it spans between sensorimotor and associative cortices, i.e., the extremes expressed in the “last-in first-out”- hypothesis.

In order to identify specific foci of cortical thinning throughout aging, we split our sample in non-overlapping groups of 5-year age intervals. We then localized thinning foci in each subgroup within the first and second functional connectivity gradient. Our results revealed that thinning foci in individual age subgroups changed along the first functional connectivity gradient (*figure 4b*). The observed trajectory across subgroups corresponds to the expected course based on the “last in-first out”-hypothesis starting from regions which mature latest in ontogeny to those maturing first: starting out in associative-transmodal cortices in youngest participants (45-50 years), the thinning focus was subsequently located closer to unimodal somatomotor brain areas in older individuals (60-65 years). In the most elderly individuals (up to 75-80 years), the thinning focus gradually “shifted back” to the center of gravity of the functional connectivity gradient manifold, specifically the first functional connectivity gradient. This either indicates a less specific, more homogeneous thinning pattern or a focus on task-related regions located in the mid of the functional network hierarchy like the ventral and dorsal attention networks. In summary, cortical thinning foci across age intervals were aligned with the principal functional connectivity gradient. Therefore, beyond its role as a principal paradigm in brain function and neurodevelopment, our results put emphasis on the relevance of the functional network hierarchy and likewise the sensorimotor association-axis in governing age-related neurodegeneration.

Regarding clinical phenotypes related to advancing age, cognitive and motor functions, such as complex information handling, executive functions, memory, mobility and overall muscle strength, are known to deteriorate over time (Bohannon and Williams Andrews, 2011; Dodds et al., 2014; Hedden and Gabrieli, 2004). We leveraged multivariate machine learning in form of a partial least squares analysis (PLS) to probe for a covariance profile relating cortical thinning and variance in phenotypical scores in the HCHS sample. We successfully identified a latent variable explaining the substantial amount of 85.10% of variance in the cortical thickness and phenotypical information from all individuals included in our study (*figure 5a*). According to this latent variable, higher cortical thickness in primary sensorimotor areas and younger age covaried with better performance in cognitive and motor assessments. Of all factors, age was identified as the most highly weighted in the phenotypical covariance pattern (*figure 5b*) and corresponded considerably with (lower) cortical thickness and lower performance in cognitive and motor tests (*figure 5d*). Of note, the found covariance construct was stable even after excluding age, sex at birth and education from the analysis indicating that there is a meaningful relationship between the remaining variables that goes beyond the interrelationship with chronological age. Taking into account the distinct correspondence with chronological age and a coherent clinical covariance profile, we hypothesize that the latent variable captures the signature of biological aging processes. In accordance with this, the thickness profile identified by PLS spatially overlapped with the map of age-related cortical thinning identified by the general linear model (*figure 5f*). Therefore, our results suggest that aging processes might lead to atrophy in the regions that contribute to good motor and cognitive function and hence late-life functionality. Coming back to our initial analysis, patterns of age-related thickness are therefore not only meaningfully contextualizable in the framework of network topology, but may also serve as a structural substrate of age-related decline in cognitive and motor functions.

The strengths of this work include the large sample size increasing statistical power for finding associations between imaging and phenotypical variables (Marek et al., 2022), the availability of high quality multimodal imaging and phenotypical data in the HCHS including a state-of-the art, robust and reproducible pipeline for image processing, and additional sensitivity analysis accounting for differing atlas parcellation schemes. Notably, the investigated age range appears particularly well suited for investigating associations between imaging markers of age-related structural brain changes and clinical phenotypes, as it captures the age interval where these clinical manifestations of aging accumulate (Buckner, 2004; Dodds et al., 2014).

Several limitations should be noted: first, our analysis is based on cross-sectional data. In comparison, a longitudinal study design can be considered superior when investigating the age-related trajectory of morphometric brain changes. Nevertheless, the identified age-related cortical thinning pattern agrees with results from existing cross-sectional and longitudinal imaging studies of a similar age range (Appleton et al., 2020; Frangou et al., 2021; Salat et al., 2004; Storsve et al., 2014; Wierenga et al., 2022). The practical constraints for conducting a longitudinal study of similar scope are considerable, both regarding human and technical resources. As a second limitation, our investigation encompasses participants aged from 45 to 80 years at time of the baseline examination, i.e., mainly the second half of the lifespan is represented. To investigate the trajectory of aging foci across the lifespan it would be of interest to investigate younger subjects as well.

## Conclusion

We identified functional and structural brain network properties linked with age-related cortical thinning in a population-based sample. Our work highlights the interplay between the connectome architecture and morphometric changes during aging with structural hubs, neighborhood alterations, and functional connectivity gradients as relevant determinants. As we found that cortical thinning occurs preferentially in regions that maintain cognitive and motor performance, our results contribute to deciphering the complex pathophysiological substrates of functional decline in older age. Collectively, our results promote the notion of age-related cortical atrophy being determined by fundamental aspects of brain network architecture.

## Materials and Methods

### Study population - the Hamburg City Health Study

Here, we investigated cross-sectional clinical and imaging data from a subgroup of the first 10,000 participants from the Hamburg City Health Study (HCHS). As described previously HCHS is an ongoing, single-center, prospective cohort study examining randomly selected citizens of the city of Hamburg, Germany, aged 45 to 74 years at time of selection (Jagodzinski et al., 2019). Participants were enrolled between 2016 and 2018 and underwent an in-depth multi-organ baseline examination with emphasis on imaging to identify risk factors, prevalence and prognostic factors for major chronic diseases. All baseline evaluations included standardized neuropsychological examinations by specifically trained medical professionals, while brain MRI was conducted in a subgroup of 2,657 participants. Hence, we analyzed data of those 2,657 participants.

### Ethics approval

The local ethics committee of the Landesärztekammer Hamburg (State of Hamburg Chamber of Medical Practitioners, PV5131) approved the study and written informed consent was obtained from all participants. Good Clinical Practice (GCP), Good Epidemiological Practice (GEP) and the Declaration of Helsinki were the ethical guidelines that governed the conduct of the study (Petersen et al., 2020).

### MRI acquisition

Images were acquired using a 3-T Siemens Skyra MRI scanner (Siemens, Erlangen, Germany). Measurements were performed with a protocol as described in previous work (Jagodzinski et al., 2019; Petersen et al., 2020). In detail, for 3D T1-weighted anatomical images, rapid acquisition gradient-echo sequence (MPRAGE) was used with the following sequence parameters: TR = 2500 ms, TE = 2.12 ms, 256 axial slices, ST = 0.94 mm, and IPR = 0.83 × 0.83 mm. Resting-state functional images were measured with the following sequence parameters: TR = 3000 ms; TE = 32 ms; flip angle = 90 degrees; FOV = 192 × 192 mm; matrix = 64×64; slices = 46; slice thickness = 3 mm; slice gap = 0 mm; effective voxel resolution = 3.0 × 3.0 × 3.0 mm. For singleshell diffusion-weighted imaging (DWI), 75 axial slices were obtained covering the whole brain with gradients (b = 1000 s/mm2) applied along 64 noncollinear directions with the following sequence parameters: repetition time (TR) = 8500 ms, echo time (TE) = 75 ms, slice thickness (ST) = 2 mm, in-plane resolution (IPR) = 2 × 2 mm, anterior–posterior phase-encoding direction, 1 b0 volume.

### Data preprocessing

For the sake of comparability and reproducibility image preprocessing was standardized based on preconfigured and containerized neuroimaging pipelines.

#### Estimation of age-related cortical thickness changes

Structural preprocessing harnessed the CAT12 surface-based morphometry pipeline (CAT12.7 r1743; https://github.com/m-wierzba/cat-container) for surface reconstruction and cortical thickness measurement building upon a projection-based thickness estimation method (Dahnke et al., 2013; Gaser et al., 2022; Rachel Aine Yotter et al., 2011; Rachel A. Yotter et al., 2011). Cortical thickness measures were normalized from individual to 32k fsLR surface space (conte69) to ensure vertex correspondence across subjects. The age-related cortical thickness alteration pattern was statistically assessed by applying a general linear model which related age and cortical thickness on a vertex-level while correcting for sex and years of education. Resulting surface maps of standardized *β* estimates of the relationship between age and cortical thickness encoded the trajectory of thickness increases (positive *β*) or reductions (negative *β*) with advancing age across all individuals. Consequently, surface maps of standardized *β* estimates were interpreted as age-related thickness alteration patterns in this work. Vertex-wise p-values were corrected for multiple comparisons based on false-discovery rate.

#### Functional and structural connectomes

Cortical thickness information derived from HCHS structural data was contextualized with group-averaged Schaefer-parcellated structural and functional connectomes from the HCHS as well as the Human Connectome Project (HCP) Young Adult dataset (Glasser et al., 2016; Schaefer et al., 2018). We opted for the Schaefer atlas as it enables assessment of results within macroscale intrinsic functional networks and facilitated sensitivity analysis across different atlas resolutions (Thomas Yeo et al., 2011). Structural HCHS connectomes were reconstructed employing QSIprep (version 0.14.2) (Cieslak et al., 2021). Group-representative structural connectomes were obtained via distance-dependent consensus thresholding (Betzel et al., 2019). Upon preprocessing of resting-state functional MRI data with fMRIPrep (version 20.2.6) HCHS functional connectomes were computed with xcpEngine (version 1.2.3) with denoising based on global signal regression and ICA-AROMA (Ciric et al., 2017; Esteban et al., 2019; Pruim et al., 2015). Negative correlations within functional connectomes were set to zero and connectomes were z-scored before group averaging. Detailed descriptions for HCHS connectome reconstructions can be found in the supplementary materials (*supplementary texts S7 and S8*). HCP connectomes are openly accessible and were downloaded as part of the ENIGMA toolbox (Larivière et al., 2021). The reconstruction approaches for the HCP connectomes have been reported previously and considerably overlap with processing choices applied to the HCHS data (Larivière et al., 2020). Put briefly, upon the HCP minimal preprocessing of structural and functional images (Glasser et al., 2016), the structural connectomes were reconstructed via application of the canonical MRtrix3 pipeline and functional connectomes were derived by calculating Pearson correlations between ROI-wise time series (Glasser et al., 2016; Tournier et al., 2019). As our work aims to draw conclusions on the age-related cortical thinning pattern from a physiologically configured brain network, we decided to report results based on the analysis of HCP-derived connectomes coming from a considerably younger sample (207 subjects, 22- 26 years) in the main analysis. Analyses based on HCHS-derived functional and structural connectomes are presented as part of the sensitivity analysis. To avoid bias by an arbitrary threshold the analysis was based on non-thresholded subject-level connectomes.

### Statistical analysis

#### Spatial correlations

To assess how the age-related cortical thickness alterations pattern corresponds with the brain connectivity profile, we related the parcel-wise *β* map to connectivity information via spatial correlations (Spearman correlation, *r_sp_*) on a group-level (*figure 2*). For every spatial correlation we performed spin permutations (n=10.000) to address the problem of spatial smoothness leading to inflated significance levels when relating two brain maps (Alexander-Bloch et al., 2018). Thereby, permutation is performed by projecting nodal information onto a sphere which is randomly rotated. After rotation information is projected back on the surface a permuted *r_sp_* is computed. Significance is assessed by relating the observed *r_sp_* to the permuted distribution.

### Connectivity profiling

Node-wise information from pre-selected concepts of functional and structural brain network connectivity were exploited to contextualize age-related cortical thickness alterations: hub ranks, neighborhood thickness alteration, and the functional network hierarchy as encoded in macroscale functional connectivity gradients. The main analysis was completely based on connectivity indices derived from Schaefer 400x7 HCP connectomes. Analysis based on alternative brain network parcellation schemes are described in the supplement.

#### Hub ranks

Hubs are commonly defined as network nodes with high intra- and intermodular connectivity. To capture both topological properties, we quantified hubs as a joint measure of weighted degree (*k*) and participation coefficient (*PC*). The degree of a node *i* was computed as the sum of all of its connection weights (Rubinov et al., 2015):

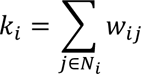

where *j* is one of the connected nodes *N_i_* and *w* is the weight of the corresponding connection.

The participation coefficient of a node *i* was calculated as:

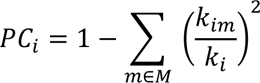

where *m* is one of the macroscale intrinsic functional networks *M*, *k_im_* is the sum of all connectivity weights between *i* and network *m* and *k* _*i*_ being the degree as described above (Thomas Yeo et al., 2011). A *PC* close to 1 signifies a node that is equivalently connected to all subnetworks of *M*, whereas a node with a value close to 0 is primarily connected to a single subnetwork. Degree and participation coefficient were ranked and combined to generate hub ranks as used for subsequent analysis.

#### Neighborhood thickness alteration

The concept of neighborhood thickness alteration (*A*) of a node *i* represents a summary measure of morphometric properties (i.e. cortical thickness) in the node neighborhood as defined by functional or structural brain network connectivity (Shafiei et al., 2020). In our study, the profile of age-related thickness alterations (*β*_*j*_) in nodes *j* connected to node *i* is averaged and weighted by their respective functional or structural seed connectivity (*w*_*ij*_):

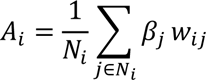

where *j* represents one of the connected nodes *N_i_*, *β*_*j*_ is its thinning profile and the corresponding connection weight *w*_*ij*_. The correction term 1/Ni is added to correct for node degree by normalizing by the number of connections. In summary, a pronounced negative *A*_*i*_ represents strong connectivity to nodes of pronounced age-related thinning.

#### Functional network hierarchy

Non-linear dimensionality reduction with diffusion map embedding of the functional connectivity matrix was performed using BrainSpace enabling localization of nodes in the functional cortical hierarchy (Margulies et al., 2016; Mesulam, 1998; Vos de Wael et al., 2020). The two components with the highest explained variance - i.e., highest eigenvalues - were propagated to further analysis as the principal and secondary functional connectivity gradient and nodewise gradient scores were extracted for each. A functional connectivity gradient can be understood as a spatial axis of connectivity variation spanning the cortical surface. Nodes of similar connectivity profiles are more closely located on these axes and nodes with less resemblance farther apart.

#### Analysis of age subintervals

In order to capture the trajectory of age-related structural brain changes we split our sample according to 5-year subintervals in non-overlapping subgroups 45-50, 50-55, 55-60, 65-70, 70-75 and 75-80 years of age. We then defined a “thinning focus” as top 10 percent of nodes with the most negative *β*. Then, to map its localization with respect to the functional network hierarchy, we calculated the median gradient scores of the thinning focus.

#### Sensitivity analysis

To assess the robustness of our results, spatial correlations, and age subgroup analyses were repeated with group-averaged connectomes derived from all investigated HCHS subjects as well as for HCP connectomes of different Schaefer atlas resolutions (100x7, 200x7; *supplementary table S3 and figure S4*).

#### Partial least squares correlation

To analyze the correspondence of cortical thickness measures with phenotypical data in individuals from the HCHS, we performed a partial least squares (PLS) correlation analysis using pyls (https://github.com/rmarkello/pyls). A detailed methodological description can be found in the supplementary materials (*supplementary figure S9 and text S10*). In brief, PLS identifies covariance profiles relating two sets of variables - here, node-wise cortical thickness (absolute values), as well as cognitive functioning (Trail Making Test A and B, Animal Naming Test, Word List Recall, Mini Mental State Exam) and motor test performances (Hand Grip Strength, Timed Up And Go Test, Chair Rising Test and Tandem Test), age, sex and years of education (Bowie and Harvey, 2006; Campagna et al., 2017; Folstein et al., 1975; Moms et al., 1989; Podsiadlo and Richardson, 1991). Cortical thickness measures were randomly permuted to assess statistical significance of latent variables. Subject-specific PLS factor scores were computed, where higher scores signify stronger expression of the identified covariance profile. Bootstrap ratios and corresponding confidence intervals were computed to quantify the node-wise contribution to the thickness-phenotypical relationship. Resulting node-wise bootstrap ratios were spatially correlated with the *β* map to probe for a potential association of covariance profiles identified by PLS and age-related cortical thickness alterations from our initial analysis.

## Acknowledgments

The authors wish to acknowledge all participants of the Hamburg City Health Study and cooperation partners, patrons and the Deanery from the University Medical Center Hamburg—Eppendorf for supporting the Hamburg City Health Study. Special thanks applies to the staff at the Epidemiological Study Center for conducting the study. The participating institutes and departments from the University Medical Center Hamburg-Eppendorf contribute all with individual and scaled budgets to the overall funding. The Hamburg City Health Study is also supported by Amgen, Astra Zeneca, Bayer, BASF, Deutsche Gesetzliche Unfallversicherung (DGUV), DIFE, the Innovative medicine initiative (IMI) under grant number No. 116074 and the Fondation Leducq under grant number 16 CVD 03., Novartis, Pfizer, Schiller, Siemens, Unilever and “Förderverein zur Förderung der HCHS e.V.”. The publication has been approved by the Steering Board of the Hamburg City Health Study.

## Author contributions

We describe contributions to the paper using the CRediT contributor role taxonomy. Conceptualization: M.P., B.C.; Data Curation: M.P., F.N., C.M., M.S, L.R.; Formal analysis: M.P.; Funding acquisition: G.T., B.C.; Investigation: M.P., F.N., C.M., M.S., L.R., E.P., S.K., J.G., U.H., J.F., R.T., C.G., G.T., B.C.; Methodology: M.P.; Software: M.P.; Supervision: G.T., B.C.; Visualization: M.P.; Writing—original draft: M.P.; Writing—review & editing: M.P., F.N., C.M., M.S., L.R., E.P., S.K., J.G., U.H., J.F., R.T., C.G., G.T., B.C.

## Data availability

Anonymized data of the analysis not published within this article will be made available on reasonable request from any qualified investigator after evaluation of the request by the Steering Board of the HCHS. Analysis code and documentation is publicly available on GitHub. A table with URLs can be found in the supplementary materials (*supplementary table S11)*.

