## Supplementary materials for "Brain Network Architecture Constrains Age-related Cortical Thinning"

### S1: Node metrics derived from Schaefer400 HCP connectomes

| Node label | FC hub rank | SC hub rank | FC neighborh. alt. | SC neighborh. alt. | FC grad. 1 | FC grad. 2 |
|---|---|---|---|---|---|---|
| 7Networks_LH_Vis_1 | 362 | 138 | -0.003887 | -0.067376 | 2.289 | 11.272 |
| 7Networks_LH_Vis_2 | 367 | 43 | -0.003155 | -0.076561 | -9.254 | 2.521 |
| 7Networks_LH_Vis_3 | 94 | 150 | -0.005784 | -0.057201 | 5.888 | 15.372 |
| 7Networks_LH_Vis_4 | 106 | 21 | -0.005592 | -0.109178 | 6.938 | 16.067 |
| 7Networks_LH_Vis_5 | 16 | 40 | -0.006527 | -0.104136 | 7.666 | 17.114 |
| 7Networks_LH_Vis_6 | 74 | 31 | -0.006210 | -0.121712 | 6.839 | 17.016 |
| 7Networks_LH_Vis_7 | 374 | 185 | -0.002427 | -0.081495 | -2.116 | 9.620 |
| 7Networks_LH_Vis_8 | 57 | 130 | -0.006070 | -0.073823 | 5.314 | 12.861 |
| 7Networks_LH_Vis_9 | 66 | 170 | -0.006368 | -0.127157 | 7.137 | 17.270 |
| 7Networks_LH_Vis_10 | 43 | 71 | -0.006064 | -0.149040 | 8.023 | 17.749 |
| 7Networks_LH_Vis_11 | 36 | 97 | -0.006290 | -0.130870 | 7.435 | 17.502 |
| 7Networks_LH_Vis_12 | 356 | 249 | -0.003126 | -0.130445 | 2.933 | 16.580 |
| 7Networks_LH_Vis_13 | 373 | 169 | -0.003268 | -0.081270 | 5.074 | 14.775 |
| 7Networks_LH_Vis_14 | 67 | 186 | -0.005443 | -0.124529 | 6.283 | 16.469 |
| 7Networks_LH_Vis_15 | 175 | 141 | -0.005634 | -0.174528 | 7.217 | 18.556 |
| 7Networks_LH_Vis_16 | 212 | 104 | -0.005342 | -0.090652 | 7.920 | 17.620 |
| 7Networks_LH_Vis_17 | 20 | 275 | -0.006510 | -0.100422 | 8.338 | 13.581 |
| 7Networks_LH_Vis_18 | 82 | 129 | -0.006430 | -0.179319 | 6.576 | 16.907 |
| 7Networks_LH_Vis_19 | 122 | 222 | -0.005127 | -0.142670 | 6.558 | 18.563 |
| 7Networks_LH_Vis_20 | 309 | 37 | -0.003402 | -0.093833 | -10.419 | 2.770 |
| 7Networks_LH_Vis_21 | 54 | 7 | -0.006705 | -0.153524 | 7.254 | 13.898 |
| 7Networks_LH_Vis_22 | 73 | 0 | -0.006577 | -0.112803 | 7.796 | 14.056 |
| 7Networks_LH_Vis_23 | 12 | 111 | -0.006460 | -0.131625 | 6.143 | 14.067 |
| 7Networks_LH_Vis_24 | 4 | 29 | -0.006794 | -0.164263 | 7.715 | 16.338 |
| 7Networks_LH_Vis_25 | 5 | 15 | -0.007025 | -0.188713 | 8.950 | 15.766 |
| 7Networks_LH_Vis_26 | 24 | 63 | -0.006942 | -0.103498 | 8.332 | 11.284 |
| 7Networks_LH_Vis_27 | 3 | 134 | -0.006584 | -0.144159 | 7.872 | 15.979 |
| 7Networks_LH_Vis_28 | 143 | 284 | -0.005848 | -0.125192 | 1.425 | 7.733 |
| 7Networks_LH_Vis_29 | 1 | 5 | -0.007214 | -0.130895 | 8.425 | 13.544 |
| 7Networks_LH_Vis_30 | 35 | 61 | -0.007024 | -0.126319 | 7.165 | 10.671 |
| 7Networks_LH_Vis_31 | 48 | 341 | -0.006176 | -0.143949 | 6.146 | 11.974 |
| 7Networks_LH_SomMot_1 | 237 | 338 | -0.005688 | -0.164603 | 11.446 | -4.260 |
| 7Networks_LH_SomMot_2 | 90 | 274 | -0.007172 | -0.150973 | 11.993 | -4.066 |
| 7Networks_LH_SomMot_3 | 310 | 179 | -0.004675 | -0.204070 | 14.958 | -4.637 |
| 7Networks_LH_SomMot_4 | 216 | 68 | -0.005339 | -0.187657 | 17.257 | -7.239 |
| 7Networks_LH_SomMot_5 | 155 | 199 | -0.005773 | -0.156902 | 16.659 | -6.052 |
| 7Networks_LH_SomMot_6 | 198 | 270 | -0.005803 | -0.175146 | 9.857 | -3.979 |
| 7Networks_LH_SomMot_7 | 166 | 232 | -0.005869 | -0.164553 | 12.609 | -3.533 |
| 7Networks_LH_SomMot_8 | 62 | 160 | -0.006262 | -0.157443 | 14.935 | -6.195 |
| 7Networks_LH_SomMot_9 | 53 | 120 | -0.006362 | -0.176608 | 13.442 | -5.420 |
| 7Networks_LH_SomMot_10 | 81 | 178 | -0.005827 | -0.170769 | 17.150 | -6.649 |

| | | | | | | |
|---|---|---|---|---|---|---|
| 7Networks_LH_SomMot_11 | 171 | 236 | -0.005474 | -0.180545 | 12.972 | -4.362 |
| 7Networks_LH_SomMot_12 | 32 | 254 | -0.006298 | -0.224348 | 17.097 | -7.847 |
| 7Networks_LH_SomMot_13 | 30 | 192 | -0.006272 | -0.258465 | 18.593 | -8.207 |
| 7Networks_LH_SomMot_14 | 55 | 229 | -0.005792 | -0.313620 | 18.566 | -8.748 |
| 7Networks_LH_SomMot_15 | 56 | 241 | -0.006212 | -0.240157 | 15.777 | -5.723 |
| 7Networks_LH_SomMot_16 | 33 | 113 | -0.006609 | -0.263292 | 15.955 | -7.157 |
| 7Networks_LH_SomMot_17 | 15 | 212 | -0.006873 | -0.348540 | 18.898 | -8.390 |
| 7Networks_LH_SomMot_18 | 168 | 95 | -0.005365 | -0.214625 | 18.456 | -6.674 |
| 7Networks_LH_SomMot_19 | 63 | 145 | -0.006134 | -0.280762 | 15.194 | -2.939 |
| 7Networks_LH_SomMot_20 | 10 | 110 | -0.006874 | -0.309383 | 17.811 | -7.374 |
| 7Networks_LH_SomMot_21 | 14 | 38 | -0.006734 | -0.247323 | 17.707 | -7.047 |
| 7Networks_LH_SomMot_22 | 95 | 220 | -0.005334 | -0.282559 | 18.415 | -8.072 |
| 7Networks_LH_SomMot_23 | 229 | 194 | -0.005023 | -0.210874 | 19.354 | -7.757 |
| 7Networks_LH_SomMot_24 | 177 | 82 | -0.004752 | -0.253655 | 20.939 | -9.412 |
| 7Networks_LH_SomMot_25 | 160 | 190 | -0.005154 | -0.208768 | 17.104 | -6.709 |
| 7Networks_LH_SomMot_26 | 41 | 4 | -0.006109 | -0.310741 | 18.602 | -7.907 |
| 7Networks_LH_SomMot_27 | 61 | 238 | -0.006310 | -0.224276 | 15.727 | -4.437 |
| 7Networks_LH_SomMot_28 | 44 | 188 | -0.005920 | -0.353242 | 19.875 | -8.666 |
| 7Networks_LH_SomMot_29 | 17 | 48 | -0.006823 | -0.269108 | 18.063 | -6.241 |
| 7Networks_LH_SomMot_30 | 244 | 221 | -0.005029 | -0.249230 | 14.497 | -3.464 |
| 7Networks_LH_SomMot_31 | 111 | 173 | -0.005598 | -0.277240 | 20.767 | -9.323 |
| 7Networks_LH_SomMot_32 | 50 | 57 | -0.006215 | -0.264468 | 19.318 | -8.019 |
| 7Networks_LH_SomMot_33 | 71 | 3 | -0.005695 | -0.314108 | 21.347 | -9.589 |
| 7Networks_LH_SomMot_34 | 156 | 13 | -0.005253 | -0.281349 | 18.376 | -7.258 |
| 7Networks_LH_SomMot_35 | 75 | 196 | -0.005752 | -0.313292 | 20.225 | -9.125 |
| 7Networks_LH_SomMot_36 | 60 | 36 | -0.005956 | -0.254902 | 18.326 | -7.758 |
| 7Networks_LH_SomMot_37 | 144 | 84 | -0.004771 | -0.285102 | 21.858 | -10.101 |
| 7Networks_LH_DorsAttn_Post_1 | 375 | 394 | -0.003689 | -0.037147 | 2.344 | 5.820 |
| 7Networks_LH_DorsAttn_Post_2 | 238 | 320 | -0.005908 | -0.089864 | -0.358 | 6.111 |
| 7Networks_LH_DorsAttn_Post_3 | 169 | 311 | -0.006238 | -0.125220 | 2.216 | 2.990 |
| 7Networks_LH_DorsAttn_Post_4 | 119 | 304 | -0.006834 | -0.098068 | 5.324 | 5.213 |
| 7Networks_LH_DorsAttn_Post_5 | 180 | 386 | -0.005240 | -0.134930 | 3.170 | 8.641 |
| 7Networks_LH_DorsAttn_Post_6 | 230 | 372 | -0.005668 | -0.211979 | 3.302 | 1.789 |
| 7Networks_LH_DorsAttn_Post_7 | 245 | 397 | -0.005525 | -0.162460 | -0.903 | 4.097 |
| 7Networks_LH_DorsAttn_Post_8 | 215 | 317 | -0.005333 | -0.226387 | 14.616 | -3.214 |
| 7Networks_LH_DorsAttn_Post_9 | 268 | 340 | -0.005379 | -0.186408 | -2.217 | 2.466 |
| 7Networks_LH_DorsAttn_Post_10 | 251 | 353 | -0.005515 | -0.145605 | 1.592 | 6.942 |
| 7Networks_LH_DorsAttn_Post_11 | 148 | 319 | -0.005834 | -0.209789 | 11.956 | -1.254 |
| 7Networks_LH_DorsAttn_Post_12 | 149 | 292 | -0.005721 | -0.205245 | 5.872 | 5.015 |
| 7Networks_LH_DorsAttn_Post_13 | 124 | 327 | -0.006211 | -0.157554 | 2.941 | 4.200 |
| 7Networks_LH_DorsAttn_Post_14 | 49 | 293 | -0.005926 | -0.154745 | 7.007 | 11.221 |
| 7Networks_LH_DorsAttn_Post_15 | 256 | 348 | -0.005722 | -0.137922 | 4.076 | 1.030 |
| 7Networks_LH_DorsAttn_Post_16 | 126 | 214 | -0.005864 | -0.236056 | 11.197 | -1.629 |

| | | | | | | |
|---|---|---|---|---|---|---|
| 7Networks_LH_DorsAttn_Post_17 | 224 | 296 | -0.005153 | -0.173095 | 15.344 | -5.144 |
| 7Networks_LH_DorsAttn_FEF_1 | 190 | 246 | -0.006290 | -0.233406 | 4.978 | 2.972 |
| 7Networks_LH_DorsAttn_FEF_2 | 323 | 257 | -0.004990 | -0.222928 | -2.621 | 2.279 |
| 7Networks_LH_DorsAttn_FEF_3 | 292 | 360 | -0.005142 | -0.208303 | 5.113 | 3.105 |
| 7Networks_LH_DorsAttn_FEF_4 | 318 | 237 | -0.005256 | -0.188768 | -1.279 | 1.172 |
| 7Networks_LH_DorsAttn_PrCv_1 | 306 | 321 | -0.005376 | -0.211718 | -1.068 | 1.858 |
| 7Networks_LH_DorsAttn_PrCv_2 | 254 | 364 | -0.005873 | -0.225973 | 1.405 | 3.669 |
| 7Networks_LH_SalVentAttn_ParOper_1 | 102 | 102 | -0.006345 | -0.191921 | 9.073 | -2.190 |
| 7Networks_LH_SalVentAttn_ParOper_2 | 335 | 379 | -0.005350 | -0.156420 | -1.751 | -1.870 |
| 7Networks_LH_SalVentAttn_ParOper_3 | 159 | 302 | -0.006157 | -0.192631 | 7.010 | -0.779 |
| 7Networks_LH_SalVentAttn_ParOper_4 | 250 | 381 | -0.005576 | -0.182457 | 1.884 | 0.299 |
| 7Networks_LH_SalVentAttn_TempOcc_1 | 263 | 208 | -0.005988 | -0.157264 | -2.672 | -0.422 |
| 7Networks_LH_SalVentAttn_FrOperIns_1 | 380 | 399 | -0.003427 | -0.150482 | 6.950 | -1.370 |
| 7Networks_LH_SalVentAttn_FrOperIns_2 | 354 | 201 | -0.004391 | -0.139745 | 6.544 | -1.561 |
| 7Networks_LH_SalVentAttn_FrOperIns_3 | 377 | 368 | -0.003877 | -0.122860 | -4.666 | -0.599 |
| 7Networks_LH_SalVentAttn_FrOperIns_4 | 357 | 166 | -0.004238 | -0.187097 | 8.400 | -2.462 |
| 7Networks_LH_SalVentAttn_FrOperIns_5 | 347 | 223 | -0.004512 | -0.168140 | 2.254 | -0.599 |
| 7Networks_LH_SalVentAttn_FrOperIns_6 | 326 | 282 | -0.004043 | -0.139660 | 7.183 | -1.973 |
| 7Networks_LH_SalVentAttn_FrOperIns_7 | 379 | 318 | -0.003993 | -0.117303 | -0.347 | -0.798 |
| 7Networks_LH_SalVentAttn_FrOperIns_8 | 142 | 180 | -0.005581 | -0.138324 | 10.201 | -3.258 |
| 7Networks_LH_SalVentAttn_FrOperIns_9 | 363 | 283 | -0.004884 | -0.175515 | -1.184 | -0.403 |
| 7Networks_LH_SalVentAttn_PFCl_1 | 282 | 35 | -0.005875 | -0.213872 | 0.658 | -0.431 |
| 7Networks_LH_SalVentAttn_Med_1 | 312 | 52 | -0.005213 | -0.112730 | 3.085 | -0.697 |
| 7Networks_LH_SalVentAttn_Med_2 | 181 | 171 | -0.005762 | -0.159037 | 13.340 | -4.399 |
| 7Networks_LH_SalVentAttn_Med_3 | 286 | 306 | -0.005610 | -0.140714 | 6.851 | 0.711 |
| 7Networks_LH_SalVentAttn_Med_4 | 225 | 216 | -0.005706 | -0.220364 | 6.852 | -0.957 |
| 7Networks_LH_SalVentAttn_Med_5 | 262 | 250 | -0.005741 | -0.116991 | 2.991 | 1.834 |
| 7Networks_LH_SalVentAttn_Med_6 | 269 | 352 | -0.005555 | -0.130844 | 5.703 | 1.694 |
| 7Networks_LH_SalVentAttn_Med_7 | 219 | 233 | -0.005393 | -0.231149 | 8.765 | -1.825 |
| 7Networks_LH_Limbic_OFC_1 | 338 | 94 | -0.000990 | -0.073056 | -16.651 | -3.297 |
| 7Networks_LH_Limbic_OFC_2 | 396 | 157 | -0.002158 | -0.074723 | -7.213 | 1.589 |
| 7Networks_LH_Limbic_OFC_3 | 341 | 83 | -0.001384 | -0.075468 | -14.058 | -2.163 |
| 7Networks_LH_Limbic_OFC_4 | 259 | 299 | -0.000915 | -0.059014 | -13.595 | -2.464 |
| 7Networks_LH_Limbic_OFC_5 | 196 | 16 | -0.003196 | -0.128097 | -15.847 | -3.914 |
| 7Networks_LH_Limbic_TempPole_1 | 378 | 90 | -0.002615 | -0.098976 | -7.064 | -1.940 |
| 7Networks_LH_Limbic_TempPole_2 | 398 | 258 | -0.001519 | -0.093786 | -2.751 | 3.510 |
| 7Networks_LH_Limbic_TempPole_3 | 389 | 248 | -0.002176 | -0.095836 | -6.400 | 2.216 |
| 7Networks_LH_Limbic_TempPole_4 | 370 | 373 | -0.001890 | -0.092494 | -12.746 | -3.028 |
| 7Networks_LH_Limbic_TempPole_5 | 385 | 242 | -0.001864 | -0.067302 | -5.995 | 1.648 |
| 7Networks_LH_Limbic_TempPole_6 | 210 | 198 | -0.003274 | -0.121400 | -12.629 | -4.394 |

| | | | | | | |
|---|---|---|---|---|---|---|
| 7Networks_LH_Limbic_TempPole_7 | 311 | 393 | -0.002419 | -0.104954 | -10.276 | -4.099 |
| 7Networks_LH_Limbic_TempPole_8 | 325 | 278 | -0.001965 | -0.112139 | -14.420 | -1.519 |
| 7Networks_LH_Cont_Par_1 | 305 | 389 | -0.004192 | -0.154938 | -9.775 | 0.755 |
| 7Networks_LH_Cont_Par_2 | 327 | 392 | -0.004454 | -0.160503 | -5.518 | -0.079 |
| 7Networks_LH_Cont_Par_3 | 131 | 347 | -0.003968 | -0.164456 | -12.468 | -1.619 |
| 7Networks_LH_Cont_Par_4 | 222 | 287 | -0.004375 | -0.168424 | -11.112 | 0.052 |
| 7Networks_LH_Cont_Par_5 | 214 | 362 | -0.004076 | -0.180211 | -10.550 | -0.282 |
| 7Networks_LH_Cont_Par_6 | 295 | 286 | -0.005047 | -0.176863 | -6.645 | 1.297 |
| 7Networks_LH_Cont_Temp_1 | 289 | 322 | -0.004099 | -0.123093 | -11.507 | -0.096 |
| 7Networks_LH_Cont_OFC_1 | 384 | 355 | -0.003010 | -0.099141 | -8.169 | -0.475 |
| 7Networks_LH_Cont_PFCl_1 | 207 | 125 | -0.003798 | -0.147126 | -12.823 | -1.756 |
| 7Networks_LH_Cont_PFCl_2 | 365 | 256 | -0.004297 | -0.179184 | -5.946 | 1.020 |
| 7Networks_LH_Cont_PFCl_3 | 345 | 235 | -0.004087 | -0.203407 | -9.587 | -0.485 |
| 7Networks_LH_Cont_PFCl_4 | 361 | 93 | -0.004087 | -0.154027 | -10.070 | -1.175 |
| 7Networks_LH_Cont_PFCl_5 | 328 | 139 | -0.005009 | -0.183558 | -3.521 | 0.340 |
| 7Networks_LH_Cont_PFCl_6 | 252 | 76 | -0.005656 | -0.199619 | -8.519 | 0.478 |
| 7Networks_LH_Cont_PFCl_7 | 358 | 398 | -0.004039 | -0.203901 | -8.850 | 0.451 |
| 7Networks_LH_Cont_PFCl_8 | 332 | 89 | -0.005085 | -0.187451 | -5.431 | -0.120 |
| 7Networks_LH_Cont_PFCv_1 | 383 | 309 | -0.002323 | -0.118367 | -7.935 | -2.058 |
| 7Networks_LH_Cont_pCun_1 | 316 | 324 | -0.004741 | -0.094975 | -8.368 | 0.200 |
| 7Networks_LH_Cont_pCun_2 | 291 | 314 | -0.005255 | -0.083999 | -8.096 | 0.446 |
| 7Networks_LH_Cont_Cing_1 | 393 | 155 | -0.002297 | -0.112868 | -4.121 | 1.839 |
| 7Networks_LH_Cont_Cing_2 | 387 | 127 | -0.002090 | -0.114990 | -12.119 | -0.983 |
| 7Networks_LH_Cont_PFCmp_1 | 313 | 59 | -0.004077 | -0.184036 | -9.466 | -0.679 |
| 7Networks_LH_Default_Temp_1 | 187 | 298 | -0.003458 | -0.138315 | -14.545 | -4.651 |
| 7Networks_LH_Default_Temp_2 | 163 | 244 | -0.003453 | -0.123616 | -15.505 | -3.825 |
| 7Networks_LH_Default_Temp_3 | 288 | 387 | -0.002918 | -0.097799 | -12.278 | -0.589 |
| 7Networks_LH_Default_Temp_4 | 107 | 135 | -0.004570 | -0.154617 | -13.938 | -3.795 |
| 7Networks_LH_Default_Temp_5 | 303 | 361 | -0.004585 | -0.191009 | 1.676 | -3.516 |
| 7Networks_LH_Default_Temp_6 | 70 | 101 | -0.004750 | -0.150420 | -14.089 | -3.897 |
| 7Networks_LH_Default_Temp_7 | 186 | 228 | -0.004631 | -0.154137 | -10.663 | -4.223 |
| 7Networks_LH_Default_Temp_8 | 261 | 330 | -0.004622 | -0.221722 | 3.289 | -3.764 |
| 7Networks_LH_Default_Temp_9 | 151 | 159 | -0.005559 | -0.180340 | 2.229 | -3.691 |
| 7Networks_LH_Default_Temp_10 | 228 | 300 | -0.004510 | -0.131105 | -7.444 | -3.763 |
| 7Networks_LH_Default_Par_1 | 202 | 316 | -0.006298 | -0.159989 | 0.289 | -0.597 |
| 7Networks_LH_Default_Par_2 | 27 | 268 | -0.005183 | -0.157644 | -14.553 | -3.736 |
| 7Networks_LH_Default_Par_3 | 179 | 230 | -0.005120 | -0.124033 | -11.154 | -0.017 |
| 7Networks_LH_Default_Par_4 | 85 | 358 | -0.004699 | -0.169974 | -16.450 | -4.079 |
| 7Networks_LH_Default_Par_5 | 47 | 182 | -0.004783 | -0.176840 | -13.979 | -3.901 |
| 7Networks_LH_Default_Par_6 | 79 | 245 | -0.004367 | -0.155439 | -15.026 | -2.456 |
| 7Networks_LH_Default_Par_7 | 113 | 285 | -0.003915 | -0.166072 | -14.269 | -2.390 |
| 7Networks_LH_Default_PFC_1 | 220 | 315 | -0.003225 | -0.101023 | -13.403 | -4.066 |
| 7Networks_LH_Default_PFC_2 | 386 | 349 | -0.002955 | -0.066544 | -9.393 | 0.728 |

| | | | | | | |
|---|---|---|---|---|---|---|
| 7Networks_LH_Default_PFC_3 | 192 | 60 | -0.003449 | -0.082928 | -16.245 | -3.439 |
| 7Networks_LH_Default_PFC_4 | 227 | 64 | -0.002552 | -0.086174 | -15.969 | -2.674 |
| 7Networks_LH_Default_PFC_5 | 200 | 346 | -0.003631 | -0.114101 | -13.746 | -4.594 |
| 7Networks_LH_Default_PFC_6 | 223 | 205 | -0.003671 | -0.148226 | -13.336 | -1.911 |
| 7Networks_LH_Default_PFC_7 | 287 | 345 | -0.003967 | -0.168382 | -10.375 | -3.817 |
| 7Networks_LH_Default_PFC_8 | 197 | 32 | -0.003036 | -0.099826 | -16.517 | -3.490 |
| 7Networks_LH_Default_PFC_9 | 204 | 19 | -0.003128 | -0.093141 | -15.237 | -2.728 |
| 7Networks_LH_Default_PFC_10 | 176 | 243 | -0.003895 | -0.162903 | -12.281 | -3.908 |
| 7Networks_LH_Default_PFC_11 | 170 | 23 | -0.003362 | -0.154926 | -15.705 | -2.207 |
| 7Networks_LH_Default_PFC_12 | 368 | 28 | -0.003660 | -0.101833 | -7.983 | -0.204 |
| 7Networks_LH_Default_PFC_13 | 99 | 11 | -0.004293 | -0.138695 | -14.804 | -4.153 |
| 7Networks_LH_Default_PFC_14 | 125 | 25 | -0.004151 | -0.210149 | -15.409 | -4.329 |
| 7Networks_LH_Default_PFC_15 | 255 | 55 | -0.003726 | -0.196142 | -13.904 | -3.679 |
| 7Networks_LH_Default_PFC_16 | 300 | 98 | -0.004216 | -0.195393 | -13.635 | -1.265 |
| 7Networks_LH_Default_PFC_17 | 83 | 165 | -0.004393 | -0.202663 | -14.641 | -3.529 |
| 7Networks_LH_Default_PFC_18 | 174 | 41 | -0.003697 | -0.202180 | -14.967 | -4.403 |
| 7Networks_LH_Default_PFC_19 | 157 | 77 | -0.003436 | -0.197307 | -16.313 | -2.978 |
| 7Networks_LH_Default_PFC_20 | 218 | 279 | -0.004646 | -0.204815 | -10.293 | -3.086 |
| 7Networks_LH_Default_PFC_21 | 154 | 147 | -0.003729 | -0.197402 | -14.328 | -2.096 |
| 7Networks_LH_Default_PFC_22 | 147 | 153 | -0.003629 | -0.194845 | -14.929 | -1.717 |
| 7Networks_LH_Default_PFC_23 | 178 | 65 | -0.003695 | -0.200434 | -13.847 | -4.079 |
| 7Networks_LH_Default_PFC_24 | 342 | 176 | -0.004025 | -0.233403 | -6.300 | -3.460 |
| 7Networks_LH_Default_pCunPCC_1 | 241 | 75 | -0.002742 | -0.072550 | -15.495 | -1.363 |
| 7Networks_LH_Default_pCunPCC_2 | 117 | 88 | -0.004173 | -0.069829 | -14.793 | -1.051 |
| 7Networks_LH_Default_pCunPCC_3 | 173 | 62 | -0.003224 | -0.055903 | -16.756 | -2.764 |
| 7Networks_LH_Default_pCunPCC_4 | 371 | 72 | -0.002050 | -0.041929 | -13.604 | -1.198 |
| 7Networks_LH_Default_pCunPCC_5 | 199 | 247 | -0.003376 | -0.059333 | -12.902 | -0.956 |
| 7Networks_LH_Default_pCunPCC_6 | 92 | 74 | -0.004221 | -0.041223 | -16.217 | -3.327 |
| 7Networks_LH_Default_pCunPCC_7 | 115 | 187 | -0.003822 | -0.037958 | -16.332 | -3.229 |
| 7Networks_LH_Default_pCunPCC_8 | 205 | 265 | -0.003231 | -0.078505 | -15.016 | -1.459 |
| 7Networks_LH_Default_pCunPCC_9 | 273 | 334 | -0.002726 | -0.086230 | -15.280 | -2.751 |
| 7Networks_LH_Default_pCunPCC_10 | 101 | 206 | -0.003994 | -0.070211 | -15.982 | -2.135 |
| 7Networks_LH_Default_pCunPCC_11 | 97 | 184 | -0.005301 | -0.064003 | -14.810 | -2.198 |
| 7Networks_RH_Vis_1 | 349 | 142 | -0.003739 | -0.081771 | 3.217 | 12.376 |
| 7Networks_RH_Vis_2 | 372 | 26 | -0.003134 | -0.084684 | -4.832 | 6.279 |
| 7Networks_RH_Vis_3 | 158 | 200 | -0.005380 | -0.038616 | 6.444 | 15.096 |
| 7Networks_RH_Vis_4 | 28 | 34 | -0.006633 | -0.100674 | 7.067 | 16.863 |
| 7Networks_RH_Vis_5 | 87 | 2 | -0.005633 | -0.106565 | 7.763 | 17.300 |
| 7Networks_RH_Vis_6 | 11 | 27 | -0.006820 | -0.118928 | 8.073 | 16.921 |
| 7Networks_RH_Vis_7 | 382 | 133 | -0.002332 | -0.103205 | 3.365 | 13.407 |
| 7Networks_RH_Vis_8 | 72 | 168 | -0.006057 | -0.117604 | 7.552 | 18.778 |
| 7Networks_RH_Vis_9 | 46 | 144 | -0.006581 | -0.109361 | 6.282 | 10.699 |
| 7Networks_RH_Vis_10 | 91 | 162 | -0.005614 | -0.128292 | 6.052 | 16.680 |

| | | | | | | |
|---|---|---|---|---|---|---|
| 7Networks_RH_Vis_11 | 23 | 154 | -0.006336 | -0.138299 | 7.173 | 16.945 |
| 7Networks_RH_Vis_12 | 84 | 167 | -0.005646 | -0.145203 | 8.694 | 17.572 |
| 7Networks_RH_Vis_13 | 352 | 143 | -0.003271 | -0.126329 | 4.042 | 16.842 |
| 7Networks_RH_Vis_14 | 275 | 44 | -0.004810 | -0.116087 | 7.357 | 17.283 |
| 7Networks_RH_Vis_15 | 100 | 18 | -0.006287 | -0.161392 | 7.235 | 18.908 |
| 7Networks_RH_Vis_16 | 38 | 344 | -0.006777 | -0.102101 | 7.491 | 8.393 |
| 7Networks_RH_Vis_17 | 344 | 271 | -0.003089 | -0.081383 | -3.938 | 8.421 |
| 7Networks_RH_Vis_18 | 37 | 118 | -0.005515 | -0.114601 | 6.878 | 17.581 |
| 7Networks_RH_Vis_19 | 42 | 39 | -0.006734 | -0.164228 | 7.469 | 15.414 |
| 7Networks_RH_Vis_20 | 52 | 1 | -0.006883 | -0.106868 | 8.209 | 12.409 |
| 7Networks_RH_Vis_21 | 247 | 217 | -0.004605 | -0.171805 | 6.589 | 19.050 |
| 7Networks_RH_Vis_22 | 8 | 121 | -0.006560 | -0.107591 | 5.690 | 13.001 |
| 7Networks_RH_Vis_23 | 7 | 53 | -0.006519 | -0.175082 | 8.379 | 18.238 |
| 7Networks_RH_Vis_24 | 22 | 45 | -0.006774 | -0.087949 | 8.411 | 11.359 |
| 7Networks_RH_Vis_25 | 9 | 50 | -0.006611 | -0.164278 | 8.889 | 17.149 |
| 7Networks_RH_Vis_26 | 0 | 17 | -0.006771 | -0.104455 | 7.234 | 15.054 |
| 7Networks_RH_Vis_27 | 31 | 384 | -0.006080 | -0.134335 | 5.375 | 13.092 |
| 7Networks_RH_Vis_28 | 19 | 47 | -0.007241 | -0.114275 | 9.396 | 9.340 |
| 7Networks_RH_Vis_29 | 2 | 9 | -0.007275 | -0.141818 | 8.267 | 11.394 |
| 7Networks_RH_Vis_30 | 29 | 259 | -0.005584 | -0.134346 | 6.100 | 13.956 |
| 7Networks_RH_SomMot_1 | 297 | 365 | -0.004789 | -0.270184 | 11.385 | -4.288 |
| 7Networks_RH_SomMot_2 | 133 | 290 | -0.005861 | -0.203845 | 7.182 | -4.098 |
| 7Networks_RH_SomMot_3 | 108 | 253 | -0.006938 | -0.191332 | 12.012 | -3.729 |
| 7Networks_RH_SomMot_4 | 321 | 122 | -0.004629 | -0.201966 | 14.081 | -4.827 |
| 7Networks_RH_SomMot_5 | 240 | 112 | -0.004628 | -0.161928 | 17.901 | -8.416 |
| 7Networks_RH_SomMot_6 | 213 | 151 | -0.005387 | -0.165337 | 17.563 | -7.640 |
| 7Networks_RH_SomMot_7 | 123 | 234 | -0.006213 | -0.209354 | 12.406 | -4.608 |
| 7Networks_RH_SomMot_8 | 140 | 326 | -0.006187 | -0.208656 | 6.885 | -0.756 |
| 7Networks_RH_SomMot_9 | 164 | 219 | -0.005340 | -0.106995 | 17.642 | -7.223 |
| 7Networks_RH_SomMot_10 | 59 | 99 | -0.006768 | -0.158126 | 15.777 | -5.993 |
| 7Networks_RH_SomMot_11 | 103 | 193 | -0.005583 | -0.096034 | 15.618 | -7.336 |
| 7Networks_RH_SomMot_12 | 68 | 137 | -0.006083 | -0.168999 | 14.302 | -5.105 |
| 7Networks_RH_SomMot_13 | 120 | 203 | -0.005521 | -0.169935 | 16.823 | -6.599 |
| 7Networks_RH_SomMot_14 | 39 | 277 | -0.006176 | -0.125085 | 16.699 | -6.739 |
| 7Networks_RH_SomMot_15 | 110 | 262 | -0.005407 | -0.213179 | 14.743 | -5.429 |
| 7Networks_RH_SomMot_16 | 6 | 128 | -0.006753 | -0.212246 | 18.310 | -8.246 |
| 7Networks_RH_SomMot_17 | 88 | 224 | -0.006071 | -0.236238 | 13.149 | -3.403 |
| 7Networks_RH_SomMot_18 | 26 | 191 | -0.006420 | -0.315664 | 18.562 | -8.488 |
| 7Networks_RH_SomMot_19 | 64 | 281 | -0.005809 | -0.239665 | 17.551 | -5.825 |
| 7Networks_RH_SomMot_20 | 283 | 337 | -0.005079 | -0.177171 | 11.087 | 1.710 |
| 7Networks_RH_SomMot_21 | 13 | 132 | -0.006722 | -0.349179 | 18.143 | -7.496 |
| 7Networks_RH_SomMot_22 | 34 | 100 | -0.006727 | -0.302753 | 12.591 | -1.512 |
| 7Networks_RH_SomMot_23 | 21 | 33 | -0.006399 | -0.295262 | 17.126 | -6.879 |

| | | | | | | |
|---|---|---|---|---|---|---|
| 7Networks_RH_SomMot_24 | 128 | 87 | -0.005272 | -0.238490 | 18.006 | -7.442 |
| 7Networks_RH_SomMot_25 | 40 | 174 | -0.006070 | -0.305566 | 18.702 | -8.010 |
| 7Networks_RH_SomMot_26 | 45 | 12 | -0.006062 | -0.327745 | 20.312 | -8.230 |
| 7Networks_RH_SomMot_27 | 25 | 67 | -0.006529 | -0.295250 | 17.721 | -5.860 |
| 7Networks_RH_SomMot_28 | 98 | 115 | -0.005903 | -0.228673 | 16.247 | -4.752 |
| 7Networks_RH_SomMot_29 | 69 | 92 | -0.005648 | -0.297350 | 19.209 | -8.541 |
| 7Networks_RH_SomMot_30 | 139 | 10 | -0.004887 | -0.289576 | 21.417 | -9.934 |
| 7Networks_RH_SomMot_31 | 231 | 156 | -0.004778 | -0.296576 | 16.117 | -4.968 |
| 7Networks_RH_SomMot_32 | 58 | 177 | -0.005835 | -0.195738 | 21.006 | -8.829 |
| 7Networks_RH_SomMot_33 | 118 | 164 | -0.005524 | -0.336332 | 21.035 | -9.235 |
| 7Networks_RH_SomMot_34 | 129 | 204 | -0.005269 | -0.268300 | 18.815 | -8.644 |
| 7Networks_RH_SomMot_35 | 51 | 114 | -0.006110 | -0.289005 | 18.611 | -7.702 |
| 7Networks_RH_SomMot_36 | 243 | 172 | -0.004324 | -0.286435 | 15.814 | -5.459 |
| 7Networks_RH_SomMot_37 | 134 | 163 | -0.005717 | -0.231527 | 15.593 | -4.498 |
| 7Networks_RH_SomMot_38 | 145 | 8 | -0.004702 | -0.302387 | 20.454 | -9.346 |
| 7Networks_RH_SomMot_39 | 132 | 14 | -0.004937 | -0.337785 | 21.919 | -9.871 |
| 7Networks_RH_SomMot_40 | 89 | 123 | -0.005494 | -0.314356 | 20.260 | -8.773 |
| 7Networks_RH_DorsAttn_Post_1 | 189 | 189 | -0.006408 | -0.103745 | 3.662 | 7.892 |
| 7Networks_RH_DorsAttn_Post_2 | 172 | 395 | -0.006522 | -0.111385 | 3.112 | 2.245 |
| 7Networks_RH_DorsAttn_Post_3 | 239 | 369 | -0.006081 | -0.150913 | 0.786 | 3.497 |
| 7Networks_RH_DorsAttn_Post_4 | 274 | 289 | -0.005057 | -0.124383 | -8.380 | 0.969 |
| 7Networks_RH_DorsAttn_Post_5 | 208 | 376 | -0.005187 | -0.113363 | 1.901 | 7.654 |
| 7Networks_RH_DorsAttn_Post_6 | 217 | 357 | -0.005179 | -0.203116 | 7.383 | 1.654 |
| 7Networks_RH_DorsAttn_Post_7 | 271 | 380 | -0.005464 | -0.115845 | -3.325 | 1.926 |
| 7Networks_RH_DorsAttn_Post_8 | 277 | 336 | -0.005083 | -0.179648 | 2.086 | 3.203 |
| 7Networks_RH_DorsAttn_Post_9 | 161 | 329 | -0.005510 | -0.222279 | 13.261 | -2.762 |
| 7Networks_RH_DorsAttn_Post_10 | 264 | 280 | -0.004971 | -0.171774 | 0.791 | 5.820 |
| 7Networks_RH_DorsAttn_Post_11 | 280 | 359 | -0.005217 | -0.176960 | -1.388 | 3.105 |
| 7Networks_RH_DorsAttn_Post_12 | 93 | 240 | -0.005972 | -0.148751 | 3.515 | 5.129 |
| 7Networks_RH_DorsAttn_Post_13 | 121 | 350 | -0.005832 | -0.243233 | 11.639 | -0.890 |
| 7Networks_RH_DorsAttn_Post_14 | 96 | 209 | -0.006071 | -0.202884 | 5.092 | 8.077 |
| 7Networks_RH_DorsAttn_Post_15 | 307 | 374 | -0.005152 | -0.080256 | -6.765 | 1.202 |
| 7Networks_RH_DorsAttn_Post_16 | 18 | 260 | -0.006356 | -0.194426 | 8.064 | 11.649 |
| 7Networks_RH_DorsAttn_Post_17 | 221 | 307 | -0.006022 | -0.119994 | 3.196 | 1.543 |
| 7Networks_RH_DorsAttn_Post_18 | 193 | 295 | -0.005820 | -0.150682 | 3.698 | 2.913 |
| 7Networks_RH_DorsAttn_Post_19 | 109 | 213 | -0.005873 | -0.214227 | 13.232 | -2.929 |
| 7Networks_RH_DorsAttn_FEF_1 | 150 | 202 | -0.006147 | -0.263754 | 4.812 | 4.520 |
| 7Networks_RH_DorsAttn_FEF_2 | 272 | 339 | -0.005150 | -0.215132 | 4.211 | 4.053 |
| 7Networks_RH_DorsAttn_FEF_3 | 298 | 288 | -0.004696 | -0.193684 | 10.133 | -0.856 |
| 7Networks_RH_DorsAttn_PrCv_1 | 278 | 266 | -0.005295 | -0.200619 | 1.606 | 2.731 |
| 7Networks_RH_SalVentAttn_TempOccPar_1 | 308 | 375 | -0.005348 | -0.162094 | -3.148 | 0.210 |
| 7Networks_RH_SalVentAttn_TempOccPar_2 | 293 | 351 | -0.005239 | -0.165648 | 2.899 | 1.425 |
| 7Networks_RH_SalVentAttn_TempOccP | 279 | 382 | -0.005593 | -0.182538 | -3.347 | -0.333 |

| | | | | | | |
|---|---|---|---|---|---|---|
| ar_3 | | | | | | |
| 7Networks_RH_SalVentAttn_TempOccPar_4 | 78 | 158 | -0.006913 | -0.142609 | 8.227 | -1.475 |
| 7Networks_RH_SalVentAttn_TempOccPar_5 | 76 | 107 | -0.006704 | -0.193839 | 8.453 | -1.784 |
| 7Networks_RH_SalVentAttn_TempOccPar_6 | 137 | 272 | -0.006322 | -0.185459 | 2.351 | 0.504 |
| 7Networks_RH_SalVentAttn_TempOccPar_7 | 294 | 378 | -0.004635 | -0.173135 | -8.604 | -0.894 |
| 7Networks_RH_SalVentAttn_PrC_1 | 258 | 251 | -0.005841 | -0.193822 | 3.114 | 1.742 |
| 7Networks_RH_SalVentAttn_FrOperIns_1 | 376 | 294 | -0.003425 | -0.077864 | 7.513 | -0.330 |
| 7Networks_RH_SalVentAttn_FrOperIns_2 | 366 | 252 | -0.003803 | -0.095783 | 4.468 | -0.899 |
| 7Networks_RH_SalVentAttn_FrOperIns_3 | 360 | 207 | -0.004027 | -0.165617 | 7.537 | -1.379 |
| 7Networks_RH_SalVentAttn_FrOperIns_4 | 319 | 161 | -0.004398 | -0.142627 | 7.933 | -1.765 |
| 7Networks_RH_SalVentAttn_FrOperIns_5 | 331 | 255 | -0.005139 | -0.121086 | -0.471 | -0.396 |
| 7Networks_RH_SalVentAttn_FrOperIns_6 | 276 | 313 | -0.004398 | -0.082423 | 9.660 | -2.941 |
| 7Networks_RH_SalVentAttn_FrOperIns_7 | 211 | 152 | -0.005479 | -0.124813 | 8.456 | -1.758 |
| 7Networks_RH_SalVentAttn_FrOperIns_8 | 322 | 333 | -0.005061 | -0.170056 | -0.704 | 0.128 |
| 7Networks_RH_SalVentAttn_PFCl_1 | 290 | 46 | -0.005706 | -0.192527 | -0.472 | -0.215 |
| 7Networks_RH_SalVentAttn_Med_1 | 265 | 108 | -0.005800 | -0.139998 | 3.866 | -0.451 |
| 7Networks_RH_SalVentAttn_Med_2 | 248 | 227 | -0.005297 | -0.149732 | 12.476 | -3.187 |
| 7Networks_RH_SalVentAttn_Med_3 | 324 | 297 | -0.005207 | -0.132435 | -0.434 | 0.113 |
| 7Networks_RH_SalVentAttn_Med_4 | 333 | 215 | -0.004992 | -0.178067 | 5.733 | 0.623 |
| 7Networks_RH_SalVentAttn_Med_5 | 235 | 291 | -0.005649 | -0.119793 | 4.978 | 4.469 |
| 7Networks_RH_SalVentAttn_Med_6 | 246 | 81 | -0.005045 | -0.200427 | 12.304 | -1.161 |
| 7Networks_RH_SalVentAttn_Med_7 | 112 | 85 | -0.005784 | -0.256311 | 11.435 | -2.706 |
| 7Networks_RH_SalVentAttn_Med_8 | 348 | 267 | -0.004403 | -0.191557 | 2.193 | -0.609 |
| 7Networks_RH_Limbic_OFC_1 | 399 | 49 | -0.000792 | -0.070651 | -5.794 | -0.775 |
| 7Networks_RH_Limbic_OFC_2 | 397 | 105 | -0.002206 | -0.061419 | -3.029 | 1.887 |
| 7Networks_RH_Limbic_OFC_3 | 232 | 119 | -0.001022 | -0.085710 | -16.255 | -3.963 |
| 7Networks_RH_Limbic_OFC_4 | 369 | 332 | -0.001626 | -0.065954 | -9.683 | -0.075 |
| 7Networks_RH_Limbic_OFC_5 | 302 | 261 | -0.000754 | -0.055178 | -12.553 | -1.792 |
| 7Networks_RH_Limbic_OFC_6 | 226 | 140 | -0.002601 | -0.103252 | -17.286 | -4.731 |
| 7Networks_RH_Limbic_TempPole_1 | 391 | 79 | -0.002422 | -0.086106 | 0.011 | 3.639 |
| 7Networks_RH_Limbic_TempPole_2 | 296 | 385 | -0.002671 | -0.097430 | -14.013 | -4.177 |
| 7Networks_RH_Limbic_TempPole_3 | 234 | 276 | -0.002691 | -0.073697 | -14.571 | -4.700 |
| 7Networks_RH_Limbic_TempPole_4 | 390 | 116 | -0.002467 | -0.088653 | -2.657 | 4.409 |
| 7Networks_RH_Limbic_TempPole_5 | 314 | 388 | -0.001687 | -0.070615 | -9.150 | -3.166 |
| 7Networks_RH_Limbic_TempPole_6 | 392 | 396 | -0.002407 | -0.066602 | 0.775 | 4.449 |
| 7Networks_RH_Limbic_TempPole_7 | 346 | 325 | -0.001908 | -0.075805 | -15.167 | -1.819 |
| 7Networks_RH_Cont_Par_1 | 146 | 263 | -0.003531 | -0.169169 | -14.209 | -2.168 |
| 7Networks_RH_Cont_Par_2 | 182 | 370 | -0.003451 | -0.189886 | -12.117 | -1.029 |
| 7Networks_RH_Cont_Par_3 | 260 | 354 | -0.004149 | -0.143571 | -10.346 | 0.396 |
| 7Networks_RH_Cont_Par_4 | 281 | 323 | -0.005079 | -0.204943 | -6.280 | 1.006 |

| | | | | | | |
|---|---|---|---|---|---|---|
| 7Networks_RH_Cont_Par_5 | 267 | 301 | -0.004126 | -0.170685 | -8.934 | 0.062 |
| 7Networks_RH_Cont_Par_6 | 194 | 269 | -0.004150 | -0.176748 | -11.462 | -0.216 |
| 7Networks_RH_Cont_Temp_1 | 233 | 390 | -0.002759 | -0.130333 | -13.428 | -1.201 |
| 7Networks_RH_Cont_Temp_2 | 270 | 273 | -0.004369 | -0.134360 | -11.137 | 0.105 |
| 7Networks_RH_Cont_PFCv_1 | 381 | 331 | -0.003403 | -0.083043 | -9.766 | -1.695 |
| 7Networks_RH_Cont_PFCl_1 | 350 | 328 | -0.002524 | -0.092659 | -11.478 | -1.112 |
| 7Networks_RH_Cont_PFCl_2 | 242 | 124 | -0.003183 | -0.114329 | -13.153 | -1.295 |
| 7Networks_RH_Cont_PFCl_3 | 185 | 78 | -0.004113 | -0.160619 | -13.373 | -1.295 |
| 7Networks_RH_Cont_PFCl_4 | 329 | 211 | -0.004729 | -0.150595 | -0.871 | 0.995 |
| 7Networks_RH_Cont_PFCl_5 | 304 | 91 | -0.004632 | -0.197331 | -7.618 | -0.217 |
| 7Networks_RH_Cont_PFCl_6 | 343 | 225 | -0.004546 | -0.185567 | -6.238 | 2.097 |
| 7Networks_RH_Cont_PFCl_7 | 336 | 181 | -0.005010 | -0.209014 | -3.335 | 0.619 |
| 7Networks_RH_Cont_PFCl_8 | 351 | 106 | -0.003586 | -0.169686 | -11.276 | -1.948 |
| 7Networks_RH_Cont_PFCl_9 | 284 | 70 | -0.004496 | -0.178888 | -10.767 | -0.102 |
| 7Networks_RH_Cont_PFCl_10 | 355 | 356 | -0.004162 | -0.184812 | -8.778 | 0.620 |
| 7Networks_RH_Cont_PFCl_11 | 334 | 96 | -0.004595 | -0.208533 | -10.297 | -0.923 |
| 7Networks_RH_Cont_PFCl_12 | 135 | 58 | -0.003623 | -0.193908 | -15.288 | -2.852 |
| 7Networks_RH_Cont_PFCl_13 | 339 | 264 | -0.004231 | -0.210709 | -10.599 | -1.737 |
| 7Networks_RH_Cont_PFCl_14 | 127 | 175 | -0.003913 | -0.197173 | -13.475 | -1.112 |
| 7Networks_RH_Cont_PFCl_15 | 340 | 239 | -0.004855 | -0.187434 | -3.341 | 1.281 |
| 7Networks_RH_Cont_pCun_1 | 317 | 308 | -0.003351 | -0.062832 | -11.272 | -0.540 |
| 7Networks_RH_Cont_pCun_2 | 104 | 183 | -0.004474 | -0.076591 | -15.526 | -1.807 |
| 7Networks_RH_Cont_Cing_1 | 388 | 149 | -0.002142 | -0.123299 | -11.254 | -0.732 |
| 7Networks_RH_Cont_Cing_2 | 395 | 195 | -0.001578 | -0.124820 | -8.964 | 0.140 |
| 7Networks_RH_Cont_PFCmp_1 | 364 | 69 | -0.003499 | -0.123171 | -10.845 | -1.249 |
| 7Networks_RH_Cont_PFCmp_2 | 320 | 54 | -0.003644 | -0.206270 | -10.737 | -0.693 |
| 7Networks_RH_Default_Par_1 | 201 | 371 | -0.005065 | -0.149771 | -10.403 | -1.856 |
| 7Networks_RH_Default_Par_2 | 162 | 377 | -0.005341 | -0.104110 | -11.004 | 0.195 |
| 7Networks_RH_Default_Par_3 | 65 | 363 | -0.004712 | -0.143288 | -15.945 | -3.648 |
| 7Networks_RH_Default_Par_4 | 77 | 383 | -0.004258 | -0.157496 | -14.160 | -2.837 |
| 7Networks_RH_Default_Par_5 | 86 | 131 | -0.004236 | -0.141772 | -15.695 | -2.216 |
| 7Networks_RH_Default_Temp_1 | 184 | 148 | -0.003621 | -0.105923 | -14.372 | -4.709 |
| 7Networks_RH_Default_Temp_2 | 165 | 335 | -0.003607 | -0.119915 | -16.565 | -4.091 |
| 7Networks_RH_Default_Temp_3 | 299 | 305 | -0.003702 | -0.136877 | -5.120 | -3.350 |
| 7Networks_RH_Default_Temp_4 | 80 | 310 | -0.004716 | -0.185527 | -10.477 | -2.965 |
| 7Networks_RH_Default_Temp_5 | 114 | 226 | -0.003880 | -0.166496 | -14.487 | -3.415 |
| 7Networks_RH_Default_Temp_6 | 337 | 303 | -0.004481 | -0.165395 | -5.609 | -2.456 |
| 7Networks_RH_Default_Temp_7 | 130 | 391 | -0.003928 | -0.118861 | -13.081 | -3.416 |
| 7Networks_RH_Default_Temp_8 | 285 | 366 | -0.004999 | -0.181094 | -5.067 | -2.751 |
| 7Networks_RH_Default_PFCv_1 | 236 | 231 | -0.002746 | -0.065799 | -14.540 | -4.151 |
| 7Networks_RH_Default_PFCv_2 | 359 | 367 | -0.003106 | -0.059085 | -12.101 | -0.182 |
| 7Networks_RH_Default_PFCv_3 | 257 | 210 | -0.004217 | -0.118840 | -11.932 | -3.536 |
| 7Networks_RH_Default_PFCv_4 | 315 | 136 | -0.004557 | -0.139945 | -10.902 | -2.921 |

| | | | | | | |
|---|---|---|---|---|---|---|
| 7Networks_RH_Default_PFCdPFCm_1 | 206 | 22 | -0.002979 | -0.093835 | -16.324 | -2.969 |
| 7Networks_RH_Default_PFCdPFCm_2 | 191 | 66 | -0.002980 | -0.126047 | -16.925 | -3.295 |
| 7Networks_RH_Default_PFCdPFCm_3 | 253 | 20 | -0.002785 | -0.094910 | -13.223 | -1.601 |
| 7Networks_RH_Default_PFCdPFCm_4 | 141 | 6 | -0.003480 | -0.115423 | -16.050 | -3.762 |
| 7Networks_RH_Default_PFCdPFCm_5 | 136 | 24 | -0.003611 | -0.174159 | -15.504 | -3.798 |
| 7Networks_RH_Default_PFCdPFCm_6 | 394 | 42 | -0.002022 | -0.107603 | -1.274 | 1.460 |
| 7Networks_RH_Default_PFCdPFCm_7 | 183 | 51 | -0.003273 | -0.195045 | -15.501 | -4.537 |
| 7Networks_RH_Default_PFCdPFCm_8 | 195 | 117 | -0.002930 | -0.182277 | -15.118 | -4.101 |
| 7Networks_RH_Default_PFCdPFCm_9 | 203 | 73 | -0.003090 | -0.182041 | -14.398 | -3.164 |
| 7Networks_RH_Default_PFCdPFCm_10 | 105 | 56 | -0.004063 | -0.177623 | -15.696 | -1.843 |
| 7Networks_RH_Default_PFCdPFCm_11 | 167 | 30 | -0.003494 | -0.187244 | -15.836 | -4.347 |
| 7Networks_RH_Default_PFCdPFCm_12 | 138 | 86 | -0.003666 | -0.177497 | -15.898 | -2.256 |
| 7Networks_RH_Default_PFCdPFCm_13 | 249 | 126 | -0.003477 | -0.175748 | -12.406 | -3.724 |
| 7Networks_RH_Default_pCunPCC_1 | 209 | 80 | -0.004073 | -0.082333 | -14.229 | -0.532 |
| 7Networks_RH_Default_pCunPCC_2 | 353 | 109 | -0.001608 | -0.051622 | -14.788 | -1.286 |
| 7Networks_RH_Default_pCunPCC_3 | 301 | 343 | -0.004217 | -0.063391 | -12.238 | -0.198 |
| 7Networks_RH_Default_pCunPCC_4 | 152 | 103 | -0.003487 | -0.067071 | -17.308 | -3.102 |
| 7Networks_RH_Default_pCunPCC_5 | 116 | 146 | -0.003757 | -0.070782 | -15.962 | -2.827 |
| 7Networks_RH_Default_pCunPCC_6 | 188 | 218 | -0.003123 | -0.070933 | -15.616 | -1.749 |
| 7Networks_RH_Default_pCunPCC_7 | 266 | 342 | -0.002426 | -0.105293 | -14.140 | -1.879 |
| 7Networks_RH_Default_pCunPCC_8 | 153 | 197 | -0.003307 | -0.072158 | -17.118 | -3.397 |
| 7Networks_RH_Default_pCunPCC_9 | 330 | 312 | -0.004544 | -0.077052 | -10.675 | -0.033 |

**S2: Scree plot for explained variance of functional gradients**

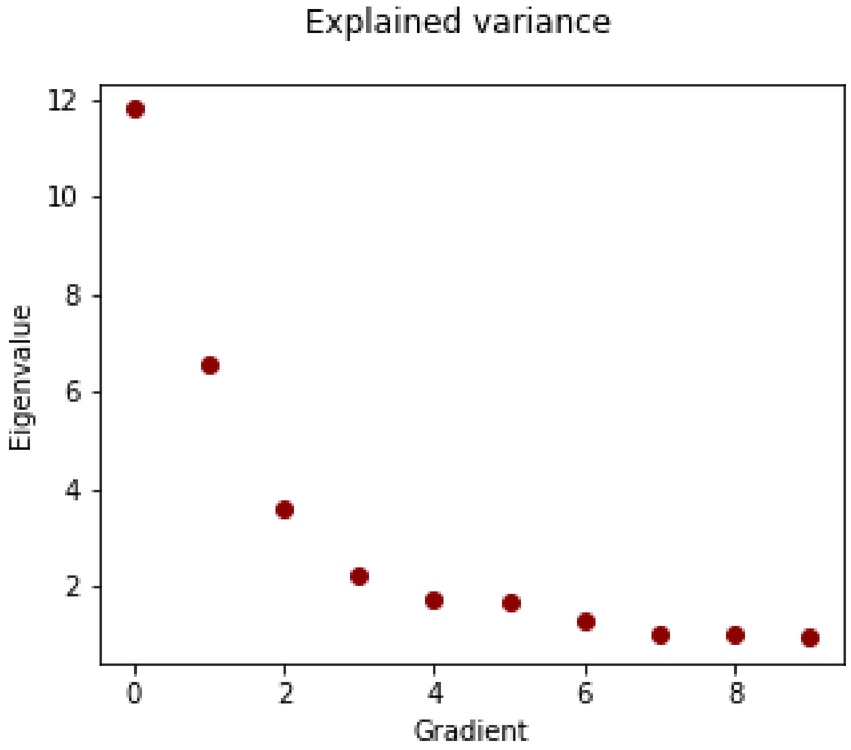

Figure S2: Scree plot of respective eigenvalues of derived functional gradients. High eigenvalues denote high explained variance.

#### S3: Sensitivity analyses

| | p Schaefer400 HCP | $r_{sp}$ Schaefer400 HCP | P Schaefer200 HCP | $r_{sp}$ Schaefer200 HCP | P Schaefer100 HCP | $r_{sp}$ Schaefer100 HCP | P Schaefer400 HCHS | $r_{sp}$ Schaefer400 HCHS |
|---|---|---|---|---|---|---|---|---|
| Functional Hubness | 0.029 | 0.27 | 0.115 | 0.22 | 0.308 | 0.11 | 0.46 | -0.02 |
| Structural Hubness | 0.047 | 0.22 | 0.037 | 0.24 | 0.004 | 0.41 | 0 | 0.26 |
| Functionally -defined Neighborh. | 0.118 | 0.23 | 0.225 | 0.16 | 0.292 | 0.13 | 0.007 | 0.41 |
| Structurally-defined Neighborh. | 0 | 0.64 | 0.003 | 0.48 | 0.033 | 0.45 | 0 | 0.44 |
| Functional Gradient 1 | 0.005 | -0.34 | 0.01 | -0.32 | 0.049 | -0.33 | 0.042 | 0.25 |
| Functional Gradient 2 | 0 | 0.44 | 0 | 0.46 | 0.408 | 0.05 | 0 | 0.47 |
| Structure-function Coupling | 0.123 | -0.16 | 0.085 | -0.17 | 0.177 | -0.1 | 0.069 | -0.25 |

#### S4: Sensitivity analyses age subintervals

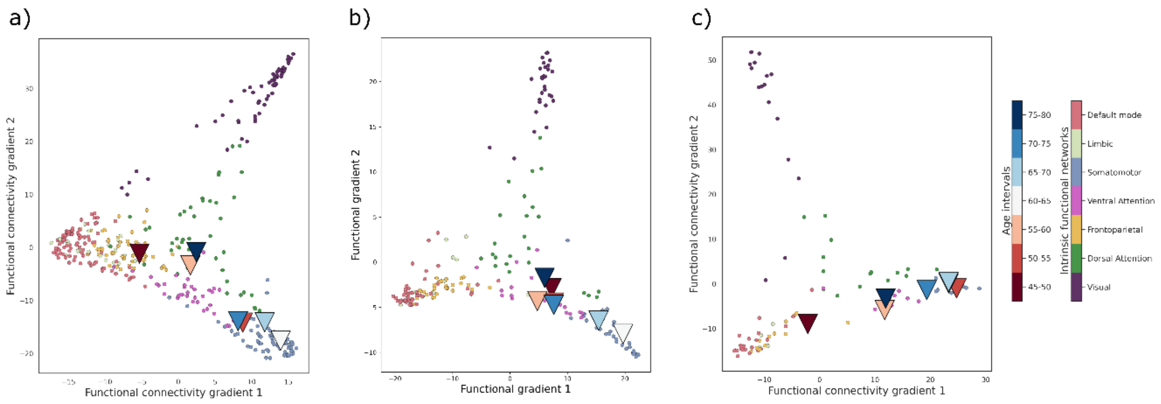

Figure S4: Sensitivity analyses of subinterval analysis. a) HCHS Schaefer400x7. b) HCP Schaefer200x7. c) HCP Schaefer100x7.

#### S5: Partial least squares analysis: latent variable statistics

| Latent variable | Explained variance (%) | p-value |
|---|---|---|
| 0 | 85.096707 | 0.000200 |
| 1 | 9.171140 | 0.000400 |
| 2 | 1.516065 | 0.039992 |
| 3 | 0.832268 | 0.188562 |
| 4 | 0.689248 | 0.729054 |
| 5 | 0.620280 | 0.434513 |
| 6 | 0.521711 | 0.733253 |
| 7 | 0.443405 | 0.677465 |
| 8 | 0.361422 | 0.926415 |
| 9 | 0.297952 | 0.972006 |
| 10 | 0.199595 | 0.668666 |
| 11 | 0.191003 | 0.999800 |
| 12 | 0.059203 | 0.956809 |

**S6: Partial least squares analysis without age, sex at birth and education**

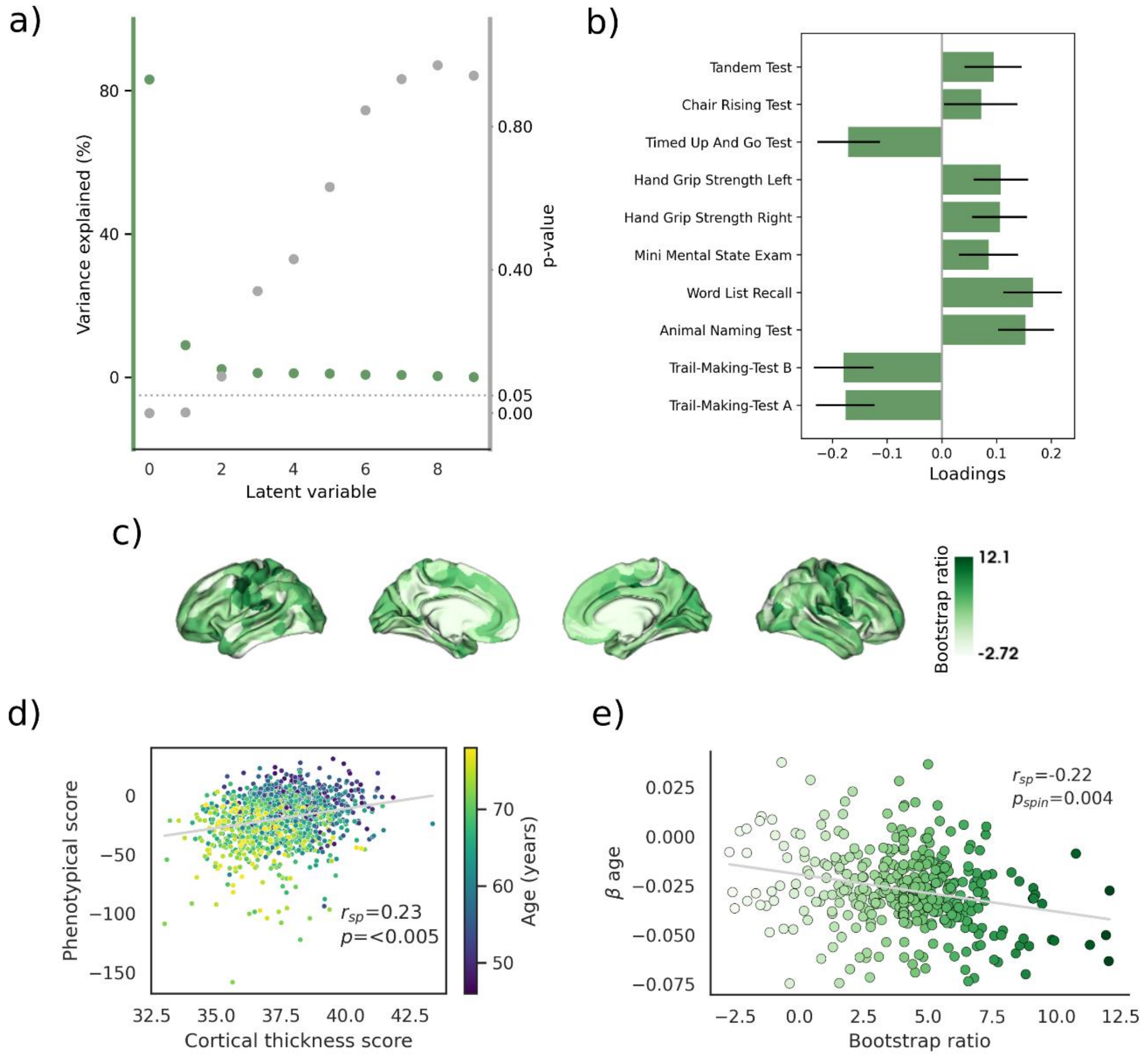

Figure S6. Partial least squares analysis results without consideration of age, sex at birth and education as phenotypical variables. Subplot order matches that in *figure 5* of the main text.

#### S7: Functional processing stream of HCHS data

Results included in this manuscript come from preprocessing performed using *fMRIPrep* 20.2.6[1,2] , which is based on *Nipype* 1.7.0[3].

**Anatomical data preprocessing**
A total of 1 T1-weighted (T1w) images were found within the input BIDS dataset.The T1-weighted (T1w) image was corrected for intensity non-uniformity (INU) with N4BiasFieldCorrection[4], distributed with ANTs 2.3.3 (https://github.com/ANTsX/ANTs), and used as T1w-reference throughout the workflow. The T1w-reference was then skull-stripped with a *Nipype* implementation of the antsBrainExtraction.sh workflow (from ANTs), using OASIS30ANTs as target template. Brain tissue segmentation of cerebrospinal fluid (CSF), white-matter (WM) and gray-matter (GM) was performed on the brain-extracted T1w using fast[5]. Brain surfaces were reconstructed using recon-all[6] [FreeSurfer 6.0.1], and the brain mask estimated previously was refined with a custom variation of the method to reconcile ANTs-derived and FreeSurfer-derived segmentations of the cortical gray-matter of Mindboggle[7]. Volume-based spatial normalization to two standard spaces (MNI152NLin6Asym, MNI152NLin2009cAsym) was performed through nonlinear registration with antsRegistration (ANTs 2.3.3), using brain-extracted versions of both T1w reference and the T1w template. The following templates were selected for spatial normalization: *FSL's MNI ICBM 152 non-linear 6th Generation Asymmetric Average Brain Stereotaxic Registration Model* [RRID:SCR_002823; TemplateFlow ID: MNI152NLin6Asym], *ICBM 152 Nonlinear Asymmetrical template version 2009c* [RRID:SCR_008796; TemplateFlow ID: MNI152NLin2009cAsym],

**Functional data preprocessing**
For each of the 1 BOLD runs found per subject (across all tasks and sessions), the following preprocessing was performed. First, a reference volume and its skull-stripped version were generated using a custom methodology of *fMRIPrep*. A deformation field to correct for susceptibility distortions was estimated based on *fMRIPrep*'s *fieldmap-less* approach. The deformation field results from co-registering the BOLD reference to the same-subject T1w-reference with its intensity inverted[8]. Registration was performed with antsRegistration (ANTs 2.3.3), and the process was regularized by constraining deformation to be nonzero only along the phase-encoding direction, and was modulated with an average fieldmap template. Based on the estimated susceptibility distortion, a corrected EPI (echo-planar imaging) reference was calculated for a more accurate co-registration with the anatomical reference. The BOLD reference was then co-registered to the T1w reference using bbregister (FreeSurfer) which implements boundary-based registration[9]. Co-registration was configured with six degrees of freedom. Head-motion parameters with respect to the BOLD reference (transformation matrices, and six corresponding rotation and translation parameters) were estimated before any spatiotemporal filtering using mcflirt[10]. BOLD runs were slice-time corrected to 1.21s (0.5 of slice acquisition range 0s-2.42s) using 3dTshift from AFNI 20160207[11]. The BOLD time-series were resampled onto the following surfaces (FreeSurfer reconstruction nomenclature): *fsnative*, *fsaverage*, *fsaverage5*. The BOLD time-series (including slice-timing correction when applied) were resampled onto their original, native space by applying a single, composite transform to correct for head-motion and

susceptibility distortions. These resampled BOLD time-series will be referred to as *preprocessed BOLD in original space*, or just *preprocessed BOLD*. The BOLD time-series were resampled into several standard spaces, correspondingly generating the following *spatially-normalized, preprocessed BOLD runs*: MNI152NLin6Asym, MNI152NLin2009cAsym. First, a reference volume and its skull-stripped version were generated using a custom methodology of *fMRIPrep*. *Grayordinates* files[12] containing 91k samples were also generated using the highest-resolution fsaverage as intermediate standardized surface space. Automatic removal of motion artifacts using independent component analysis[13] was performed on the *preprocessed BOLD on MNI space* time-series after removal of non-steady state volumes and spatial smoothing with an isotropic, Gaussian kernel of 6mm FWHM (full-width half-maximum). Corresponding "non-aggressively" denoised runs were produced after such smoothing. Additionally, the "aggressive" noise-regressors were collected and placed in the corresponding confounds file. Several confounding time-series were calculated based on the *preprocessed BOLD*: framewise displacement (FD), DVARS and three region-wise global signals. FD was computed using two formulations following Power (absolute sum of relative motions[14]) and Jenkinson (relative root mean square displacement between affines[10]). FD and DVARS were calculated for each functional run, both using their implementations in *Nipype* [following the definitions by [14]]. The three global signals were extracted within the CSF, the WM, and the whole-brain masks. Additionally, a set of physiological regressors were extracted to allow for component-based noise correction [CompCor[15]]. Principal components were estimated after high-pass filtering the *preprocessed BOLD* time-series (using a discrete cosine filter with 128s cut-off) for the two *CompCor* variants: temporal (tCompCor) and anatomical (aCompCor). tCompCor components were then calculated from the top 2% variable voxels within the brain mask. For aCompCor, three probabilistic masks (CSF, WM and combined CSF+WM) were generated in anatomical space. The implementation differs from that of Behzadi et al. in that instead of eroding the masks by 2 pixels on BOLD space, the aCompCor masks were subtracted by a mask of pixels that contains a volume fraction of GM. This mask was obtained by dilating a GM mask extracted from the FreeSurfer's *aseg* segmentation, and it ensured that components were not extracted from voxels containing a minimal fraction of GM. Finally, these masks were resampled into BOLD space and binarized by thresholding at 0.99 (as in the original implementation). Components were also calculated separately within the WM and CSF masks. For each CompCor decomposition, the *k* components with the largest singular values were retained, so that the retained components' time series were sufficient to explain 50 percent of variance across the nuisance mask (CSF, WM, combined, or temporal). The remaining components were dropped from consideration. The head-motion estimates calculated in the correction step were also placed within the corresponding confounds file. The confound time series derived from head motion estimates and global signals were expanded with the inclusion of temporal derivatives and quadratic terms for each[16]. Frames that exceeded a threshold of 0.5 mm FD or 1.5 standardised DVARS were annotated as motion outliers. All resamplings could be performed with *a single interpolation step* by composing all the pertinent transformations (i.e. head-motion transform matrices, susceptibility distortion correction when available, and co-registrations to anatomical and output spaces). Gridded (volumetric) resamplings were performed using antsApplyTransforms (ANTs), configured with Lanczos interpolation to minimize the smoothing effects of other kernels. Non-gridded (surface) resamplings were performed using mri_vol2surf (FreeSurfer).

Many internal operations of *fMRIPrep* use *Nilearn*[17] 0.6.2, mostly within the functional processing workflow. For more details of the pipeline, see the section corresponding to workflows in *fMRIPrep*'s documentation.

Copyright waiver

The above boilerplate text was automatically generated by fMRIPrep with the express intention that users should copy and paste this text into their manuscripts *unchanged*. It is released under the CC0 license.

#### S8: Structural processing stream of HCHS data

Preprocessing was performed using *QSIPrep* 0.14.2, which is based on *Nipype* 1.6.1[3].

##### Anatomical data preprocessing

The T1-weighted (T1w) image was corrected for intensity non-uniformity (INU) using N4BiasFieldCorrection[4] [ANTs 2.3.1], and used as T1w-reference throughout the workflow. The T1w-reference was then skull-stripped using antsBrainExtraction.sh (ANTs 2.3.1), using OASIS as target template. Spatial normalization to the ICBM 152 Nonlinear Asymmetrical template version 2009c was performed through nonlinear registration with antsRegistration [ANTs 2.3.1, https://github.com/ANTsX/ANTs], using brain-extracted versions of both T1w volume and template. Brain tissue segmentation of cerebrospinal fluid (CSF), white-matter (WM) and gray-matter (GM) was performed on the brain-extracted T1w using FAST[5] [FSL 6.0.3:b862cdd5].

##### Diffusion data preprocessing

Any images with a b-value less than 100 s/mm^2 were treated as a *b*=0 image. MP-PCA denoising as implemented in MRtrix3's dwidenoise[18] was applied with a 5-voxel window. After MP-PCA, Gibbs unringing was performed using MRtrix3's mrdegibbs[19]. Following unringing, B1 field inhomogeneity was corrected using dwibiascorrect from MRtrix3 with the N4 algorithm[4]. After B1 bias correction, the mean intensity of the DWI series was adjusted so all the mean intensity of the b=0 images matched across each separate DWI scanning sequence.

FSL (version 6.0.3:b862cdd5)'s eddy was used for head motion correction and Eddy current correction[20]. Eddy was configured with a *q*-space smoothing factor of 10, a total of 5 iterations, and 1000 voxels used to estimate hyperparameters. A linear first level model and a linear second level model were used to characterize Eddy current-related spatial distortion. *q*-space coordinates were forcefully assigned to shells. Field offset was attempted to be separated from subject movement. Shells were aligned post-eddy. Eddy's outlier replacement was run. Data were grouped by slice, only including values from slices determined to contain at least 250 intracerebral voxels. Groups deviating by more than 4 standard deviations from the prediction had their data replaced with imputed values. Final interpolation was performed using the jac method.

A deformation field to correct for susceptibility distortions was estimated based on *fmriprep*'s *fieldmap-less* approach. The deformation field is resulting from co-registering the b0 reference to the same-subject T1w-reference with its intensity inverted[8]. Registration was performed with antsRegistration (ANTs 2.3.1), and the process regularized by constraining deformation to be nonzero only along the phase-encoding direction, and modulated with an average fieldmap template. Based on the estimated susceptibility distortion, an unwarped b=0 reference was calculated for a more accurate co-registration with the anatomical reference. Several confounding time-series were calculated based on the preprocessed DWI: framewise displacement (FD) using the implementation in *Nipype* [following the definitions by [14]]. The head-motion estimates calculated in the correction step were also placed within the corresponding confounds file. Slicewise cross correlation was also calculated. The DWI time-series were resampled to ACPC, generating a *preprocessed DWI run in ACPC space* with 2mm isotropic voxels.

**MRtrix3 Reconstruction**

Multi-tissue fiber response functions were estimated using the dhollander algorithm. FODs were estimated via constrained spherical deconvolution[21] (CSD) using an unsupervised multi-tissue method[22,23]. A single-shell-optimized multi-tissue CSD was performed using MRtrix3Tissue (https://3Tissue.github.io), a fork of MRtrix3[24] FODs were intensity-normalized using mtnormalize[25].

Many internal operations of *QSIPrep* use *Nilearn*[17] and *Dipy*[26]. For more details of the pipeline, see the section corresponding to workflows in *QSIPrep*'s documentation.

#### S9: Illustration partial least squares analysis

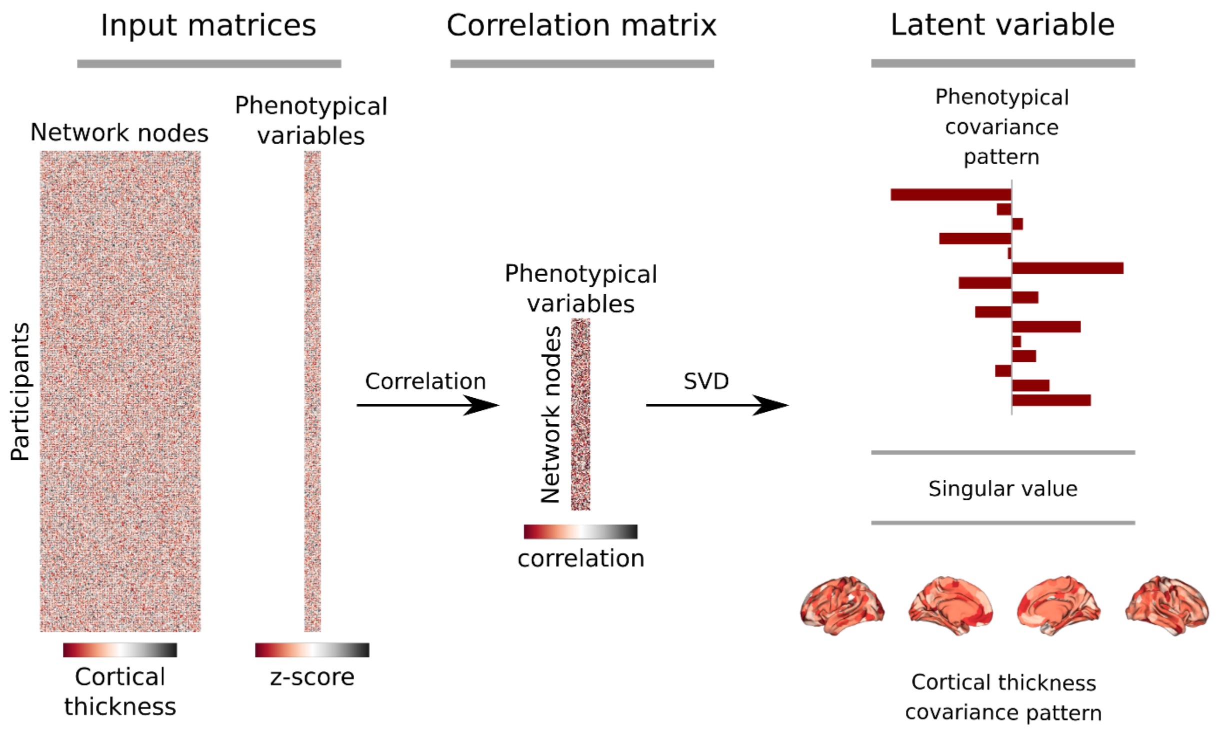

Figure S9: Partial least squares methodology. Please refer to S5 for a detailed description of the approach. This figure is a modification from figure 1 of Zeighami et al., 2019[27].

#### S10: Detailed explanation partial least squares analysis

Thickness and phenotypical data were arranged in two matrices $X_{n_{participants} \times n_{nodes}}$ and $Y_{n_{participants} \times n_{phenotypical\ variables}}$, z-scored and the thickness-phenotypical correlation matrix was then calculated which encompassed the correlation of all nodal cortical thickness values and phenotypical measures across subjects. Upon that singular value decomposition was performed on the correlation matrix resulting in a set of mutually orthogonal latent variables (LVs). The number of latent variables is determined by the smaller dimension of the covariance matrix - i.e., its rank; here the number of phenotypical variables. A latent variable consists of a left singular vector, a right singular vector and a singular value. Singular value decomposition results in matrices of left ($U_{n_{nodes} \times n_{phenotypical\ variables}}$) and right ($V_{n_{phenotypical\ variables} \times n_{phenotypical\ variables}}$) singular vectors as well as a diagonal matrix of singular values ($\Delta_{n_{phenotypical\ variables} \times n_{phenotypical\ variables}}$). A singular vector weights the corresponding original variables so that their covariance is maximized. The weighted nodal values can be understood as maximally covarying thickness patterns and their corresponding manifestation phenotypical data. Each pairing of left and right singular vector is connected by a singular value which is propotional to the covariance of both. The explained covariance of a latent variable was assessed as the ratio of the squared singular value to the sum of the other squared singular values. Significance of a latent variable was assessed by comparing the observed explained variance to a non-parametric distribution of permuted values via permutation (n=5000) of the subject-order in $X$. To measure to which extent a covariance pattern represented by a latent variable is manifested within a subject, participant-specific scores were calculated by projecting $U$ on $X$ for a thickness score

$$Thickness\ score = UX$$

and $V$ on $Y$ for a phenotypical score

$$Phenotypical\ score = VY$$

which can be thought of as a component or factor weighting in principal component analysis or factor analysis, respectively. Bootstrap resampling was used to identify nodes with a considerable contribution to the thickness-phenotypical relationship. For that, participants were randomly sampled with replacement from $X$ and $Y$ (n=5000) which results in a set of resampled covariance matrices propagated to singular value decomposition. This yields a sampling distribution for each node enabling the computation of a bootstrap ratio as the ratio of its singular vector weight and the corresponding standard error as estimated from bootstrapping. The bootstrap ratio is used to measure a node's contribution to the observed covariance pattern represented by a latent variable as relevant nodes exhibit a high singular vector weight and have a small standard error which means they are stable across subjects. The bootstrap ratio pattern was spatially correlated with the thickness $\beta$-map to assess whether the age-related cortical thinning pattern corresponds with the nodes contributing to the covariance pattern.

#### S11: Code availability

| Analysis step | *Code URL* |
|---|---|
| DWI preprocessing with QSIPrep | https://github.com/csi-hamburg/CSIframe/blob/709275c816b7746bf7168f69b652b2aec569b838/pipelines/qsiprep/qsiprep.sh |
| fMRI preprocessing with fMRIPrep | https://github.com/csi-hamburg/CSIframe/blob/main/pipelines/fmriprep/fmriprep.sh |
| Functional connectome reconstruction | https://github.com/csi-hamburg/CSIframe/blob/main/pipelines/xcpengine/xcpengine.sh |
| Spatial correlations, partial least squares analysis and age subinterval analysis | https://github.com/csi-hamburg/2022_petersen_aging_ct_connectivity |
| Structural processing with CAT | https://github.com/csi-hamburg/CSIframe/blob/709275c816b7746bf7168f69b652b2aec569b838/pipelines/cat12/cat12.sh |
